# T6SS-mediated competition by *Stenotrophomonas rhizophila* shapes seed-borne bacterial communities and seed-to-seedling transmission dynamics

**DOI:** 10.1101/2024.07.22.604635

**Authors:** Tiffany Garin, Agathe Brault, Coralie Marais, Martial Briand, Anne Préveaux, Sophie Bonneau, Marie Simonin, Matthieu Barret, Alain Sarniguet

**Author notes:** **Corresponding author:** Alain Sarniguet.

## Abstract

Seeds harbor diverse microbial communities important for plant growth and health. During germination, seed exudation triggers intense microbial competition, shaping the communities transmitted to seedlings. This study explores the role of the bacterial type VI secretion system (T6SS)-mediated interference competition in seed microbiota transmission to seedlings. The analysis of T6SS distribution within 180 genome sequences of seed-borne bacterial strains enabled the construction of synthetic communities (SynCom) with different levels of phylogenetic diversity and T6SS richness. These SynComs were inoculated with *Stenotrophomonas rhizophila* CFBP13503, a bacterial strain possessing an active T6SS *in vitro* and *in planta*. The impact of T6SS on SynCom composition was assessed *in vitro* by comparing CFBP13503 wild-type strain or its isogenic T6SS-deficient mutant co-inoculation. Additionally, the effects of T6SS on bacterial community dynamics during seed-to-seedling transmission were examined following seed-inoculation. The T6SS of *S. rhizophila* CFBP13503 targets a broad range of bacteria belonging to 5 different orders. The susceptibility of competing bacteria was partly explained by their phylogenetic proximity and metabolic overlap with CFBP13503. Furthermore, the T6SS modulates the relative abundance of specific bacterial taxa during seed-to-seedling transmission, depending on the initial seed inoculum and plant developmental stage. Depending on the sensitivity of the co-inoculated competitors, the T6SS can provide a competitive advantage to CFBP13503, resulting in an increase in population size.

**Importance:** The high prevalence of T6SS in seed-borne bacteria supports the importance of T6SS-mediated competition for seed microbiota assembly. *In vitro*, *S. rhizophila* CFBP13503 T6SS exerts a strong impact on bacterial community dynamics. The susceptibility to T6SS increases with the phylogenetic and metabolic proximity of bacteria to CFBP13503, suggesting an influence of interspecies trophic patterns in T6SS-mediated competitions. *In planta* and in soil, CFBP13503 T6SS influences specific bacterial taxa, leading to shifts in bacterial interactions and distinct community dynamics. T6SS-mediated competition plays a pivotal role in shaping seed bacterial communities and the dynamics of seed-to-seedling transitions.

## INTRODUCTION

Seeds serve as a vector for the dissemination of numerous plant-associated microorganisms (1). While seeds are often considered as the starting point for the assembly of plant microbiota (2), the percentage of seed-borne microorganisms successfully transmitted to seedlings is quite variable (3–7). These variations in seedling transmission of seed-borne microbes may be attributed to competition with soil-borne taxa for seedling colonization (8). This competition between microorganisms is the result of the exudation of numerous molecules during seed imbibition, which creates a specific habitat - the spermosphere (9). Like other habitats (i.e. rhizosphere and phyllosphere), the nature and quantities of exudates is an important driver of microbial community composition (10). This is perhaps even more striking within the spermosphere since the plant defense responses appear to be not induced during germination (11, 12). Hence microbial competition is probably a key factor for successful seedling colonization.

Microbial competition could be broadly divided into exploitative and interference competitions (13). The former is based on the rapid and efficient use of resources, a strategy used by copiotroph (14). High relative abundance of copiotrophic bacterial taxa have been reported in seedlings (15, 16) and spermospheres (10) of different plant species. These observations suggested that this life history strategy is important for seed to seedling transmission. In contrast to exploitative competition, interference competition relies on limiting the access of other microorganisms to resources through the production of antimicrobial compounds (13). Among the wide range of mechanisms used to inhibit microbial growth, contact-dependent mechanisms based on the Type VI Secretion System (T6SS) have been well studied (17).

T6SS is a contractile nanomachine found in many gram-negative bacteria. T6SSs are categorized into four subtypes: T6SS^i^ mainly found in Pseudomonadota and subdivided into subgroups i1 to i5 (18–21), T6SS^ii^ specific to *Francisella* (22, 23), T6SS^iii^ present in Bacteroidota (24), and T6SS^iv^ found in *Amoebophilus asiaticus* (25). T6SS is used by bacteria to inject effectors directly into prokaryotic or eukaryotic cells and, for few, in proximal environments (26). Toxic effectors are associated with the arrow of the system composed of the Hcp tube capped with the VgrG/PAAR spike. Surrounded by a contractile sheath, the arrow is propelled by mechanical force, allowing the delivery of toxic effectors into the target cell. Effectors are associated with cognate immunity proteins to prevent cell self-intoxication (18, 27). Other non-immune defense mechanisms can also provide protection against T6SS attack, such as spatial separation and rapid growth, general stress responses, and the presence of physical barriers preventing T6SS from reaching its target (28, 29).

Competition-relatedness theory postulates that competition should be more intense between closely related species that may share similar functions and substrate affinities. While this has been demonstrated in interbacterial antagonism experiments involving diffusible compounds (30), it remains uncertain whether this concept applies to contact-dependent mechanisms like T6SS. This is especially relevant in the context of seed-associated bacteria, where competition in the spermosphere plays a crucial role in seedling colonization. In this environment, where both seed-borne and soil-borne microbes compete for space and resources, understanding T6SS-mediated competition could offer insights into which bacteria are more likely to establish in the seedling microbiota. Therefore, we aimed to investigate whether competition-relatedness theory can be extended to explain T6SS-mediated interactions within seed-associated communities. To investigate this theory extension, we rely on the strong T6SS-mediated antibiosis provided by a *Stenotrophomonas rhizophila* strain (CFBP13503) *in vitro* against the phytopathogenic bacterium *Xanthomonas campestris* pv. *campestris* 8004 (Xcc8004) (31). *S. rhizophila* CFBP13503 is a seed-borne bacterial strain isolated from radish (32) that displays a high transmission rate from seed to seedling (33) and limits Xcc8004 populations *in planta* in a T6SS-manner (31). In the present work, the wild-type strain of *S. rhizophila* CFBP13503 and its isogenic T6SS-deficient mutant were inoculated in synthetic communities (SynComs) of different phylogenetic diversity in a culture medium. If competition-relatedness theory applies to our study model, then we anticipate a greater effect of CFBP13503 T6SS in communities with low phylogenetic diversity compared to those with high phylogenetic diversity. This experimental system also enabled the estimation of the range of bacterial competitors targeted by the *S. rhizophila* T6SS *in vitro*. Finally, we deployed the SynCom approach *in planta* for analyzing the significance of S. *rhizophila* T6SS during seed to seedling transmission.

## MATERIALS AND METHODS

### Bacterial strain collection from seed and genome sequencing

Bacterial isolates were collected from seed lots of radish (*Raphanus sativus*), winter oil seed rape (*Brassica napus*) and common bean (*Phaseolus vulgaris*) (**Table S1**). For each seed lot, 500 to 1,000 seeds were soaked in 4 ml of PBS supplemented with Tween20 (0.05% v/v), for 2h30 (radish and rapeseed) and 16h (bean) at 4°C under constant agitation. Suspensions were serially diluted and plated on Tryptic Soy Agar 1/10 (17 g.L^-1^ tryptone, 3 g.L^-1^ soy peptone, 2.5 g.L^-1^ glucose, 5 g.L^-1^ NaCl, 5 g.L^-1^ K2HPO4 and 15 g.L^-1^ agar) supplemented with cycloheximide (50 µg.mL^-1^). After 7 days incubation at 18°C, 200 to 300 colonies per seed lot were subcultured from several Petri dishes of different colony density (10 to 150) into 96-well plates containing 200 µL Tryptic Soy Broth. After 5 days of incubation at 18°C with constant agitation (70 rpm), 10 µL of bacterial suspension were transferred to 90 µL of sterile water for subsequent PCR amplification. 190 µL sterile 80% glycerol were added to each well of the remaining bacterial suspensions and the plates were stored at -80°C. Molecular typing of each bacterial isolate was conducted with a portion of the *gyrB* gene following the procedure described by Armanhi et al. (2016) (34). For the detailed procedure also applied to bacterial community metabarcoding see the “*gyrB* amplification sequencing” section.

A total of 180 bacterial isolates were subjected to Illumina HiSeq 4000 PE150 sequencing. These isolates were chosen by comparing their *gyrB* haplotypes with *gyrB* sequences previously obtained on seed samples. All genome sequences were assembled with SOAPdenovo 2.04 (35) and VELVET 1.2 (36).

### Phylogenetic analysis, prediction of functional trait and assessment of T6SS distribution

Phylogenomics of the 180 bacterial strains isolated from seeds and seedlings of *B. napus*, *P. vulgaris* and *R. sativus* (**Table S1**) was performed using the following procedure (37). Briefly, single-copy genes detected in all genome sequences were extracted with AMPHORA2 (38). Twenty-four single-copy genes were selected for subsequent phylogenetic analysis. Sequence alignment was performed with MUSCLE. Alignments were concatenated and a maximum likelihood phylogenetic tree was built with RAxML v8 (39). Phylogenetic tree was displayed and annotated with iTol v6.8 (40). The pairwise phylogenetic distances between *S. rhizophila* CFBP13503 and each bacterial strain was computed with the cophenetic.phylo function of the ape v5.7.1 package (41).

Prediction of functional traits from bacterial genomes was performed with microTrait v1.0.0 (42). Resource acquisition traits (at granularity 3) were selected. The resource overlap between CFBP13503 and bacterial strains was calculated by dividing the number of shared resource traits (between CFBP13503 and each strain) by the total number of resource traits of CFBP13503.

Detection of T6SS within genome sequences is based on the search for 10 Clusters of Orthologous Groups (COG3515, COG3516, COG3517, COG3518, COG3519, COG3520, COG3521, COG3522, COG3455, COG3523) specifics to T6SS (43). Classification of T6SS was performed through analysis of TssB protein sequences (COG3516) with the SecReT6 database v3 (44).

### *In vitro* SynCom confrontation assays

Forty-five bacterial strains isolated from seeds of radish and rapeseed were used to build SynComs (**Table S2, Fig. 1A** and **Fig. 1B**). These strains possess unique *gyrB* Amplicon Sequence Variants (ASVs) and are representative of the range of variable prevalence in seed microbiota (**Fig. 1b**). Five SynComs with different phylogenetic diversity and number of T6SS loci were constructed (**Fig. 1C** and **Fig. 1D**). Each of these five SynComs were supplemented with Xcc8004, a strain previously identified as sensitive to the T6SS of *S. rhizophila* CFBP13503 (31). Finally, each SynCom was mixed with either the strain *S. rhizophila* CFBP13503 or the isogenic T6SS deficient-mutant Δ*hcp* (31).

**Figure 1.**
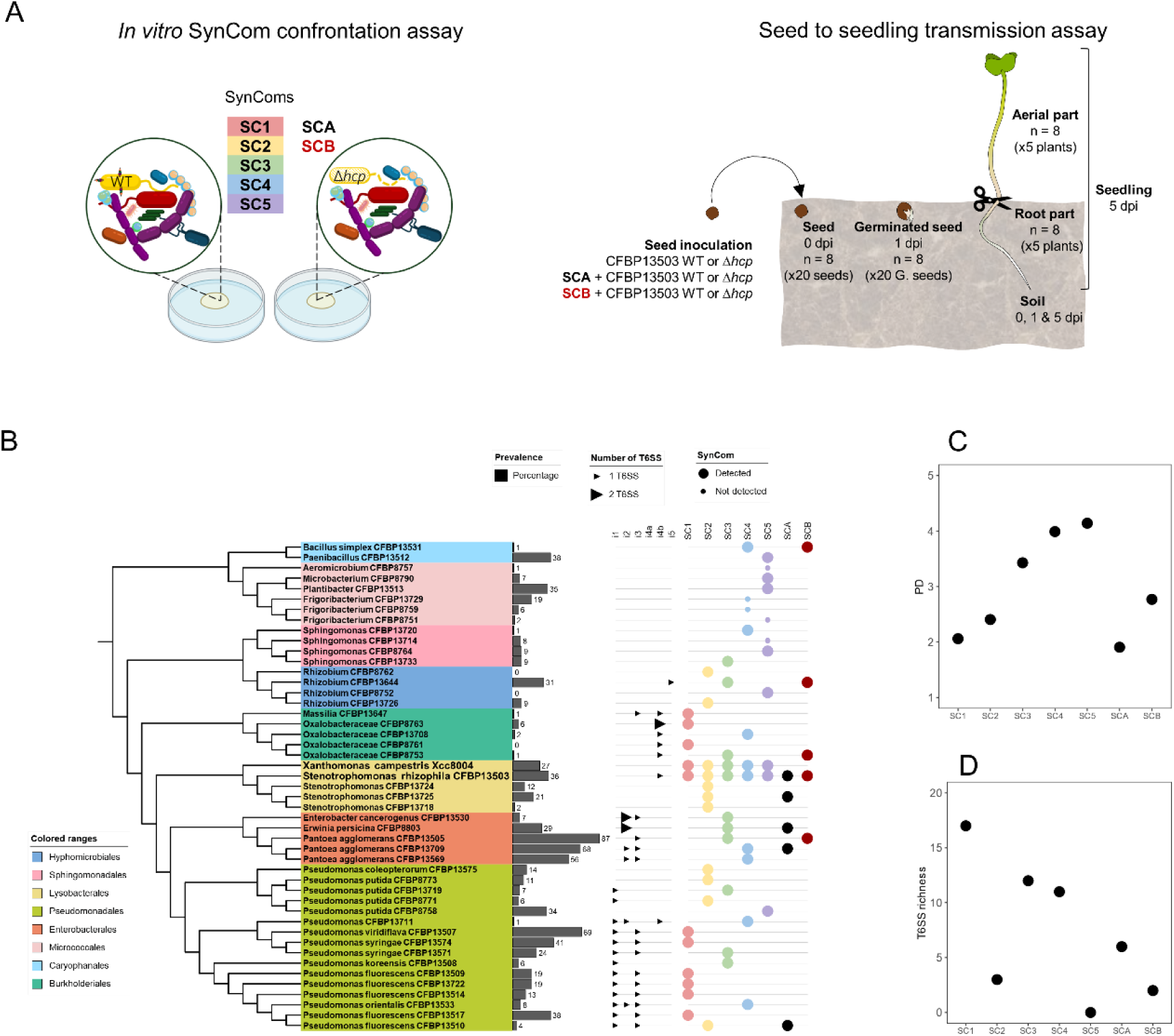
Overview of experimental designs and strain sensitivity to T6SS. The impact of the T6SS of *S. rhizophila* CFBP13503 on the composition of synthetic communities (SynCom; SC) was assessed *in vitro* across five SynComs (SC1 to SC5) and *in planta* across two SynComs (SCA and SCB) (**A**). The phylogenetic relationships of the selected bacterial strains, the prevalence in 740 seed samples (1), the associated T6SS clusters and the distribution in SynComs are depicted in (**B**). SynComs were designed with different phylogenetic diversity (**C**) and T6SS richness (**D**).

To construct each SynCom, bacterial strains were cultured overnight in 10 ml of TSB10% at 28°C under constant agitation (150 rpm). Overnight cultures were centrifuged (4,000 g, 8 min) and the resulting pellets were resuspended in sterile water at a bacterial concentration of 5.10^8^ cells.ml^-1^. Bacterial concentrations were determined according to correspondence between OD_600nm_ and number of colony-forming units (CFUs). SynCom were obtained by mixing equivalent volumes of each bacterial suspension (n=10). These initial SynCom mixtures were then combined with equal volumes of *S. rhizophila* CFBP13503 or the isogenic T6SS deficient-mutant, resulting in confrontation mixtures containing 50% of CFBP13503 and 50% of the other SynCom members. Twenty µl drops (n=10) were spotted on TSA10% and incubated at 28°C for 6 and 24 hours (4 repetitions per SynCom). Simultaneously, 500 µl of each initial confrontation mixture were centrifuged (11,000 g, 5 min) and the resulting pellets were stored at -80°C for subsequent DNA extraction. After 6 and 24 hours of confrontation, inoculated spots were resuspended in 1.5 ml and 5 ml of sterile water, respectively. 500 µl of these cell suspensions were centrifuged (11,000 g, 5 min) and the pellets were stored at -80°C. DNA extraction was performed with the Wizard® Genomic DNA Purification Kit (Promega) using supplier recommendations.

The relative abundance of each bacterial strain was assessed using *gyrB* amplicon sequencing (see below). To evaluate the enrichment or depletion of bacterial strains in response to the T6SS of *S. rhizophila* CFBP13503, we use the following formula log_2_[(RA_6h_/RA_0h_)_WT_ / (RA_6h_/RA_0h_)_Δhcp_], which measures change in relative abundance (RA) between the initial (0h) and 6-hour time points (6h) in presence (WT) and absence (Δ*hcp*) of functioning T6SS.

Total bacterial abundance in all the SynComs of the *in vitro* confrontation experiment was quantified by qPCR. DNA samples used for previous metabarcoding experiments were amplified with the primers Com1 (CAGCAGCCGCGGTAATAC) and 769R (ATCCTGTTTGMTMCCCVCRC) targeting a 270 bp region from the bacterial 16S rRNA gene (45). qPCR was performed in duplicates in a 15 µl reaction mix containing 7.5 µl of 2X SsoAdvanced™ Universal SYBR® Green Supermix (Bio-Rad), 0.3 µl of each primer (10 µM), 2 µl of DNA template and 4.9 µl of pure water, with a CFX96 Touch™ Cycler (Bio-Rad). Cycling conditions consisted of an initial denaturation step (5 min, 95°C), then 40 cycles of 10 sec at 95°C and 22 sec at 59°C. A reference target range was achieved with 10^-3^ to 10^-8^ serial dilutions of the CFBP13503 amplicon target. The melting curve was determined after the last amplification cycle using a temperature range of 59 to 95°C and a temperature transition rate of 0.5°C. The Cq values and the melting temperature of amplification products were calculated automatically. The reaction efficiency percentage was calculated using serial dilutions with the formula: E% = (10(−1/slope)-1)/100 and ranged from 70 to 100%. The Cq values were converted to copy numbers of gene targets per nanogram of DNA, using standard curves generated from amplicon targets as a reference and the values were normalized to represent the average copy number of targets per microliter according to the initial volume of sample used for DNA extraction. Changes in 16S rRNA gene copy number were considered as significant at a p-value < 0.05 (Wilcoxon test).

### Pairwise confrontation assay *in vitro*

Change in strain abundance following confrontation with CFBP13503 WT or the isogenic T6SS-deficient mutant was also measured by pairwise confrontation assays. Twenty phylogenetically distinct strains were selected. Spontaneous rifampicin-resistant (Rif^R^) variants were generated for each strain. Variants with no growth differences (TSB10%, 24h, 500 rpm) from the wild-type strains were selected.

*S. rhizophila* CFBP13503, CFBP13503Δ*hcp* and the selected Rif^R^ bacterial strains were cultured overnight in 10 mL of TSB10% (28°C, 150 rpm). Overnight cultures were centrifuged (4,000 g, 8 min) and the resulting pellets were resuspended in sterile water at an OD_600nm_ of 0.5. One hundred µl of Rif^R^ bacterial suspension and 100 µl of *S. rhizophila* CFBP13503 (WT or Δ*hcp*) suspensions were mixed. As a control, 100 µl of bacterial suspensions were mixed with 100 µL of sterile water. Three individual 20 µl drops were deposited on TSA10% and incubated at 28°C for 6 hours and 24 hours. Inoculated spots were resuspended in 2.5 ml of sterile water, serial diluted and plated on TSA10% either containing spectinomycin (50 µg.ml^-1^) and ampicillin (100 µg.ml^-1^) for *S. rhizophila* or rifampicin (50 µg.ml^-1^) for the Rif^R^ bacterial strains. Changes in CFU abundances were considered as significant at a p-value < 0.05 (Wilcoxon test). The detection limit was 1 CFU/plate at the most concentrated suspension. All data below this detection threshold,10^3^ CFU/ individual seedling, were removed.

### Impact of the T6SS of CFBP13503 on bacterial community compositions during seedling transmission

The impact of *S. rhizophila* CFBP13503 T6SS on seedling transmission was evaluated by inoculating sterilized radish seeds (*Raphanus sativus* var. Flamboyant5) with *S. rhizophila* CFBP13503 WT or the isogenic T6SS-deficient mutant. Both strains were inoculated on radish seeds alone or in the presence of two SynComs, SCA and SCB (**Fig. 1A** and **Fig. 1B**). SCA was composed of 4 T6SS-sensitive strains (CFBP8803, CFBP13510, CFBP13709 and CFBP13725), whereas SCB was assembled with 2 T6SS-sensitive strains (CFBP8753 and CFBP13505) and 2 T6SS-resistant strains (CFBP13531 and CFBP13644). Xcc8004 was not added in SCA and SCB because of irregular detection when introduced in the different SynComs tested *in vitro*. Therefore, we chose other resistant or sensitive strains efficiently tracked in SynComs. The relative abundance of each strain was measured using *gyrB* amplicon sequencing on 8 individual seeds just before sowing (0 dpi), 8 individual germinated seeds after sowing (1 dpi) and 8 individual seedlings (5 dpi).

Bacterial suspensions were prepared at a concentration of 10^9^ cells.ml^-1^ from 24h TSA10% plates. SynComs were obtained by mixing equivalent volumes of each bacterial suspension (n=4). These initial SynCom mixtures were then combined with equal volumes of *S. rhizophila* CFBP13503 or the isogenic T6SS deficient-mutant, resulting in mixtures containing 50% of *S. rhizophila* and 50% of the other SynCom members. In addition, both *S. rhizophila* strains (WT or Δ*hcp*) were inoculated alone on seeds at a concentration of 10^9^ cells.ml^-1^ mixed with an equivalent volume of sterile water.

Radish seeds were surface-sterilized using the protocol employed in Simonin et al., 2023 (33). Briefly, seeds were sonicated (40 Hertz, 1 min), then immersed in 96° ethanol (1 min), 2.6% sodium hypochlorite (5 min), 96° ethanol (30 s) and rinsed three times with sterile water. Sterilized seeds were dried (30 min) on sterile paper and distributed in subsamples of 60 seeds. These subsamples were inoculated with: sterile water (uninoculated control), CFBP13503WT, CFBP13503*Δhcp,* SCA+CFBP1503WT, SCA+CFBP13503*Δhcp*, SCB+CFBP13503WT, SCB+CFBP13503Δ*hcp* and incubated at 20°C (70 rpm, 15 min) before being dried on sterile paper (15 min). Forty seeds from each condition were subsequently sown in seedling cell trays containing unsterile, pre-moistened potting soil (Traysubstrat 75/25, Klasmann-Deilmann, Germany) at a depth of 5mm. The trays were then placed in a growth chamber (photoperiod: 16h/8h, 20°C). Eight remaining seeds were individually resuspended in 1.5 ml of sterile water by vortexing for 30 seconds. Five hundred µl were stored at -80°C and 200 µl were diluted and plated on TSA10% (total bacteria) and TSA10%+Amp100+Spe50 (selective medium for CFBP13503). Eight germinated seeds (1 dpi) per condition were individually placed in sterile bags containing 2 ml of sterile water and 500 µl and 200 µl of the resulting suspensions were stored at -80°C and diluted and plated, respectively. Eight seedlings per condition were collected at 5 dpi. The aerial parts and the roots were separated, removing as much adhering soil as possible. Each part was placed individually in sterile bags with 2 mL of sterile water and processed like germinated seeds. At each time point (0, 1 and 5 dpi) soil samples were collected and stored at -80°C.

### *gyrB* amplicon sequencing

Community profiling was performed through *gyrB* amplicon sequencing following the procedure described earlier (15). Briefly the first PCR step was performed with the gyrB_aF64/gyrB_aR553 primer set using the AccuPrimeTM Taq DNA Polymerase (Invitrogen, Carlsbad, California, USA). PCR products were purified with Sera-MagTM (Merck, Kenilworth, New Jersey). Illumina adapters and barcodes were added with a second PCR. PCR products were purified and pooled at equimolar concentration. The pooled amplicon library was quantified with the KAPA Library Quantification Kit (Roche, Basel, Switzerland). Amplicon library was mixed with 5% of PhiX and sequenced with four MiSeq reagent kits v3 600 cycles (Illumina, San Diego, California, USA). A blank extraction kit control, a PCR-negative control and PCR-positive control (*Lactococcus piscium* DSM 6634, a fish pathogen that is not plant-associated) were included in each PCR plate. The raw amplicon sequencing data are available on the Sequence Read Archive (SRA) with the accession number PRJNA1127244.

Sequence analyses of *gyrB* were performed as follows. Primer sequences were removed with cutadapt 2.7 (46) and trimmed fastq files were processed with DADA2 v 1.10 (47). The following parameters were employed for sequence filtering and trimming: maxN=0, maxEE=c(2,2), truncQ=3. The truncLen parameter was adapted for each sequencing run (25th percentile < Q30). Chimeric sequences were removed with the removeBimeraDenovo function of DADA2. Post-clustering curation of Amplicon Sequence Variant (ASV) was performed with LULU v 0.1.0 (48) at a similarity threshold of 99%. Taxonomic affiliation of each ASVs was performed with a naive Bayesian classifier (49) with our in-house *gyrB* database (train_set_gyrB_v5.fa.gz) available upon request. Unassigned sequences at the phylum level and *parE* sequences (a *gyrB* paralog) were filtered. The identification of sequence contaminants was assessed using decontam (50) v 1.8.0 at a threshold of 0.3.

Diversity analyses were conducted with PhyloSeq 1.40.0 (51). Alpha-diversity indexes (richness and Pielou’s evenness) were calculated after rarefaction of 2,700 (*in vitro* assay) and 5,000 (*in planta* assay) reads per sample. Changes in bacterial phylogenetic composition were estimated with weighted UniFrac distances (52). A permutational multivariate analysis of variance was performed with the adonis2 function of vegan 2.6.4. Changes in relative abundance was assessed with the microViz v 0.10.5 package (53) using a beta-binomial regression model provided in the corncob v 0.3.1 package (54). Taxa were filtered at a minimum prevalence threshold of 0.2. Changes in relative abundances were considered as significant at a p-value < 0.05.

## RESULTS

### T6SS prevalence among seed-associated bacteria

In order to build SynComs with different levels of T6SS diversity, we sequenced 180 bacterial genomes isolated from radish, bean and rapeseed seeds and seedlings (**Table S1**). Among these 180 genomes 61.7% (111 genomes) possess at least one T6SS cluster (**Fig. 2**). T6SS is detected in all bacterial orders analyzed, except for genomes of monoderms (e.g. *Caryophanales, Micrococcales* and *Propionibacteriales*), which do not contain diderm-like protein secretion systems (54). Most T6SS clusters (n=229) belong to T6SS subtype i (n=226), while subtype iii was restricted to the Bacteroidota order *Flavobacteriales* (n=3). Among T6SS subtype i, the following groups were detected: i1 (n=57), i2 (n=63), i3 (n=80), i4a (n=1), i4b (n=21), i5 (n=4). Seed-associated bacteria possess on average 1.6 T6SS clusters per genome with a range of 1 to 5 clusters per genome. Overall, T6SS is highly prevalent among seed-associated bacteria and might play a significant role for the colonization of the seed habitat.

**Figure 2.**
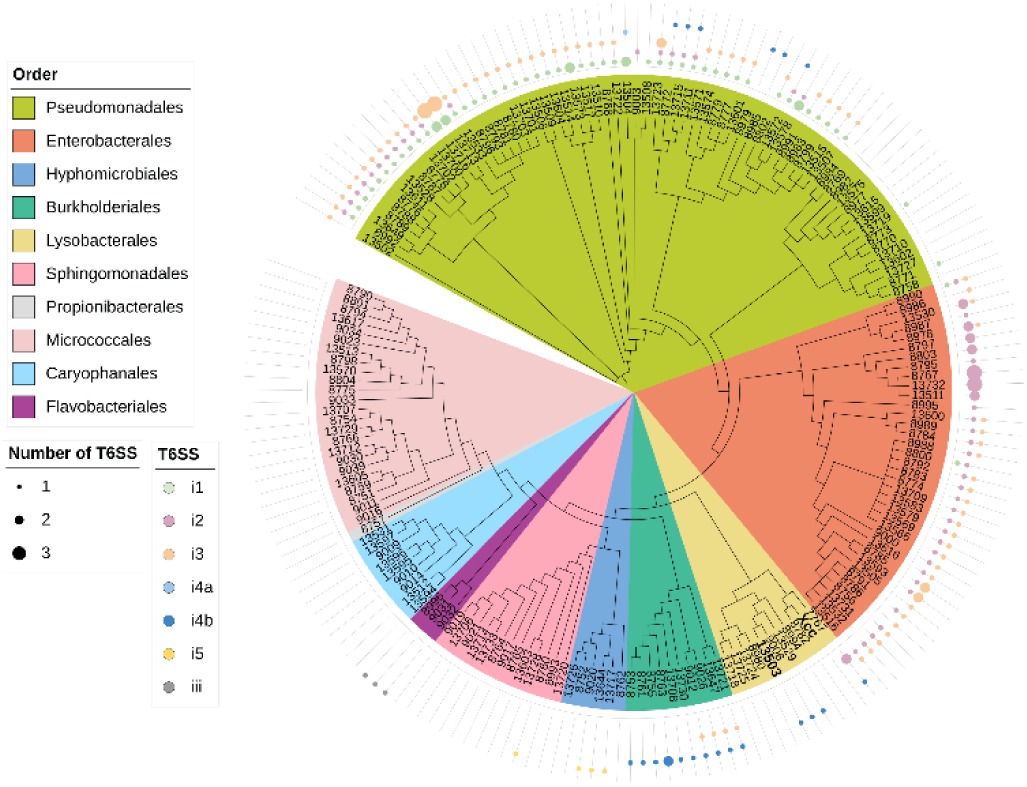
T6SS distribution among seed-associated bacteria. Maximum likelihood (PhyML) phylogenetic tree was constructed using 24 marker genes from a dataset of 180 genomes of seed-associated bacterial strains of rapeseed, radish and bean. Bacteria are color-coded according to their phylogenetic orders. T6SS clusters are colored according to their classification, which was determined through an analysis of the conserved TssB protein. The sizes of the shapes correspond to the number of T6SS belonging to a specific group present in the bacterial genomes.

### *S. rhizophila* CFBP13503 T6SS shapes the structure of bacterial SynComs *in vitro*

To analyze the impact of a T6SS secretion system on the assembly of bacterial communities associated with seeds, we chose a strain of *S. rhizophila*, CFBP13503, whose T6SS is capable of inhibiting the growth of *X. campestris* pv. *campestris* (Xcc) under *in vitro* conditions (31). We constructed 5 SynComs with contrasting levels of observed phylogenetic diversity (Faith’s PD, **Fig. 1C**) and T6SS richness (**Fig. 1D**). These SynComs were either inoculated with the wild-type (WT) strain CFBP13503 or with an isogenic Δ*hcp* mutant, in which the deletion of the Hcp component prevents the T6SS from functioning properly, leading to a loss of competitive killing ability (31).

According to 16S rRNA gene copy number, population sizes of all SynComs were significantly reduced by the T6SS of *S. rhizophila* CFBP13503 after 6h of confrontation (**Fig. 3A**). After 24h of confrontation, the difference in population size between SynComs inoculated with the WT or T6SS-deficient strains disappeared or decreased in amplitude (**Fig. 3A**).

**Figure 3.**
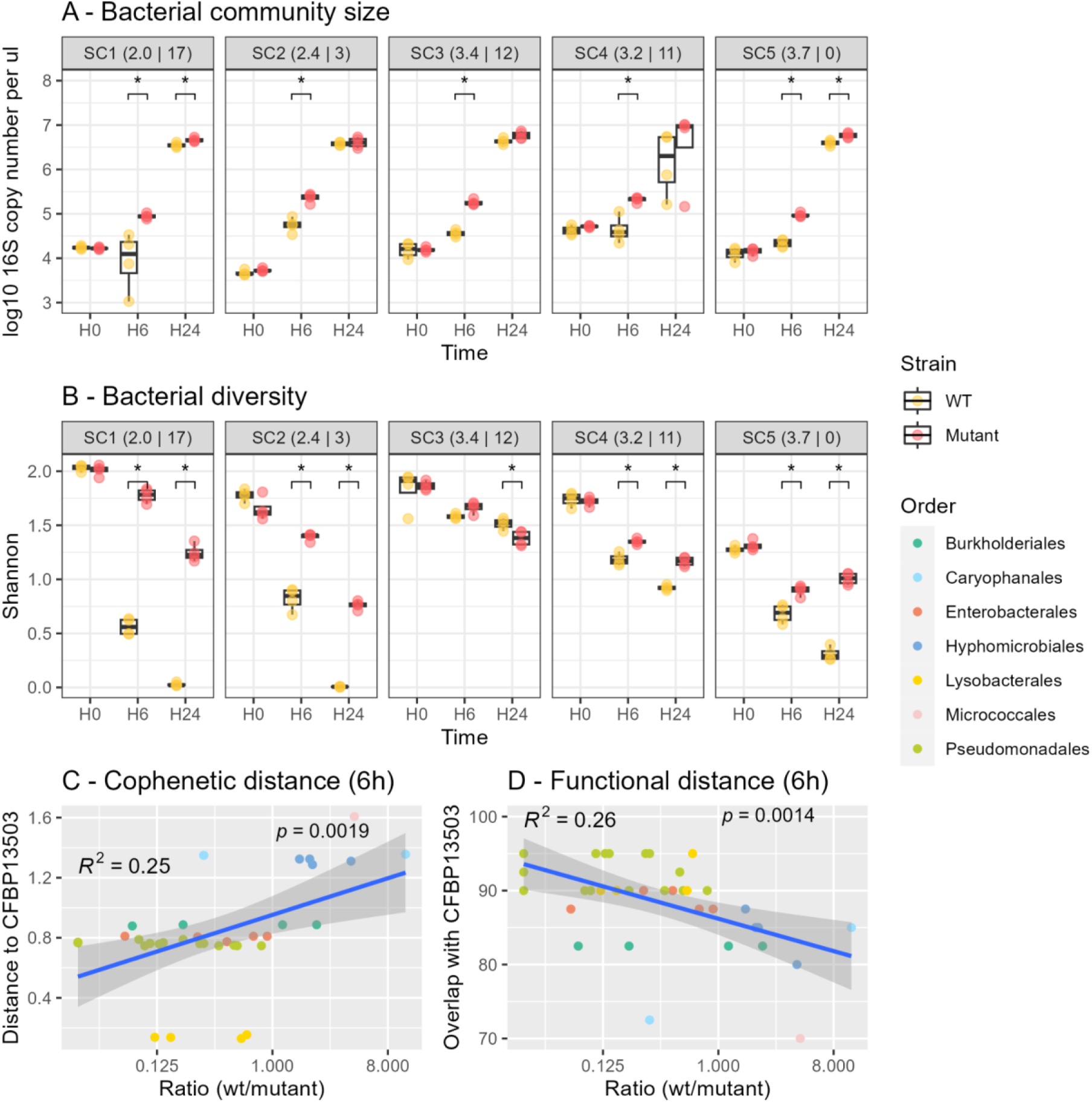
T6SS of *S. rhizophila* CFBP13503 modulated SynCom diversity. Bacterial community size (**A**) and Shannon diversity (**B**) of bacterial communities during confrontations with *S. rhizophila* CFBP13503 wild type (WT) and the isogenic T6SS mutant Δ*hcp* at different time points (0h, 6h and 24h). The phylogenetic diversity (Faith’s index) and number of T6SS clusters per SynCom are indicated in parentheses. Correlation between changes in relative abundance of SynCom members at 6h and their cophenetic distances to CFBP13503 (**C**). Correlation between changes in relative abundance of SynCom members at 6h and their resource overlap with CFBP13503 (**D**).

SynComs composition was then estimated by *gyrB* amplicon sequencing. Of the 3,233,153 reads obtained after sequence processing, 99.5% corresponded to the *gyrB* sequences of the inoculated strains. In terms of sensitivity, 42 strains out of the 47 inoculated were detected in at least one time point. The strains *Microbacterium* CFBP8790, *Frigoribacterium* CFBP13729, CFBP8759, CFBP8751, and *Sphingomonas* CFBP13714 that were not detected in our *gyrB* metabarcoding analysis were therefore excluded from the final data analysis.

Confrontations between SynComs and CFBP13503 strains (WT or Δ*hcp*) significantly (*P <* 0.001) impacted the phylogenetic composition (weighted UniFrac distance; PERMANOVA, P<0.0001) of all SynComs, explaining 26% to 75% of variance (**Table S3**). A significant increase (*P <* 0.05) in Shannon diversity index was observed from time 6h for almost all SynComs (with the exception of SC3) when communities were inoculated with Δ*hcp* (**Fig. 3B**). The magnitude of this increase in alpha-diversity was greater at 24 h, except once again for SC3. Phylogenetic diversity of SynCom alone did not explain changes in Shannon diversity since SynComs with equivalent levels of phylogenetic (e.g. SC3 and SC5) responded differently to the T6SS of *S. rhizophila* CFBP13503 (**Fig. 3B**).

We hypothesized that communities with a higher T6SS-content would be more resistant to the T6SS of CFBP13503 because of potential killing of CFPB13503 or higher resistance to CFBP13503 effectors *via* the presence of a greater diversity of immunity proteins possessed by members of these communities. This would then result in smaller modulations of community size and diversity. According to qPCR (**Fig. 3A**) and metabarcoding (**Fig. 3B**) this is clearly not the case.

Community composition changes were further analyzed by tracking relative abundance shifts of SynCom members (**Fig. S1**). A decrease in relative abundance of strains affiliated to *Pseudomonadales*, *Enterobacterales*, and *Lysobacterales* was observed during confrontation with the WT (**Fig. 4A**). In contrast, relative abundance of strains belonging to *Caryophanales*, *Hyphomicrobiales* and *Micrococcales* were enriched when inoculated with the WT (**Fig. 4A**). Members of these bacterial orders responded uniformly to the WT strain with consistent decrease or increase in relative abundance **(Fig. 4A)**. However, the response within Burkholderiales was more variable, with some strains showing a decrease in relative abundance and others exhibiting an increase in the presence of the WT strain **(Fig. 4A).** Overall, these results suggest that there is a T6SS-dependent shift in the relative abundance of phylogenetically diverse taxa in SynComs when confronted with *S. rhizophila* CFBP13503.

**Figure 4.**
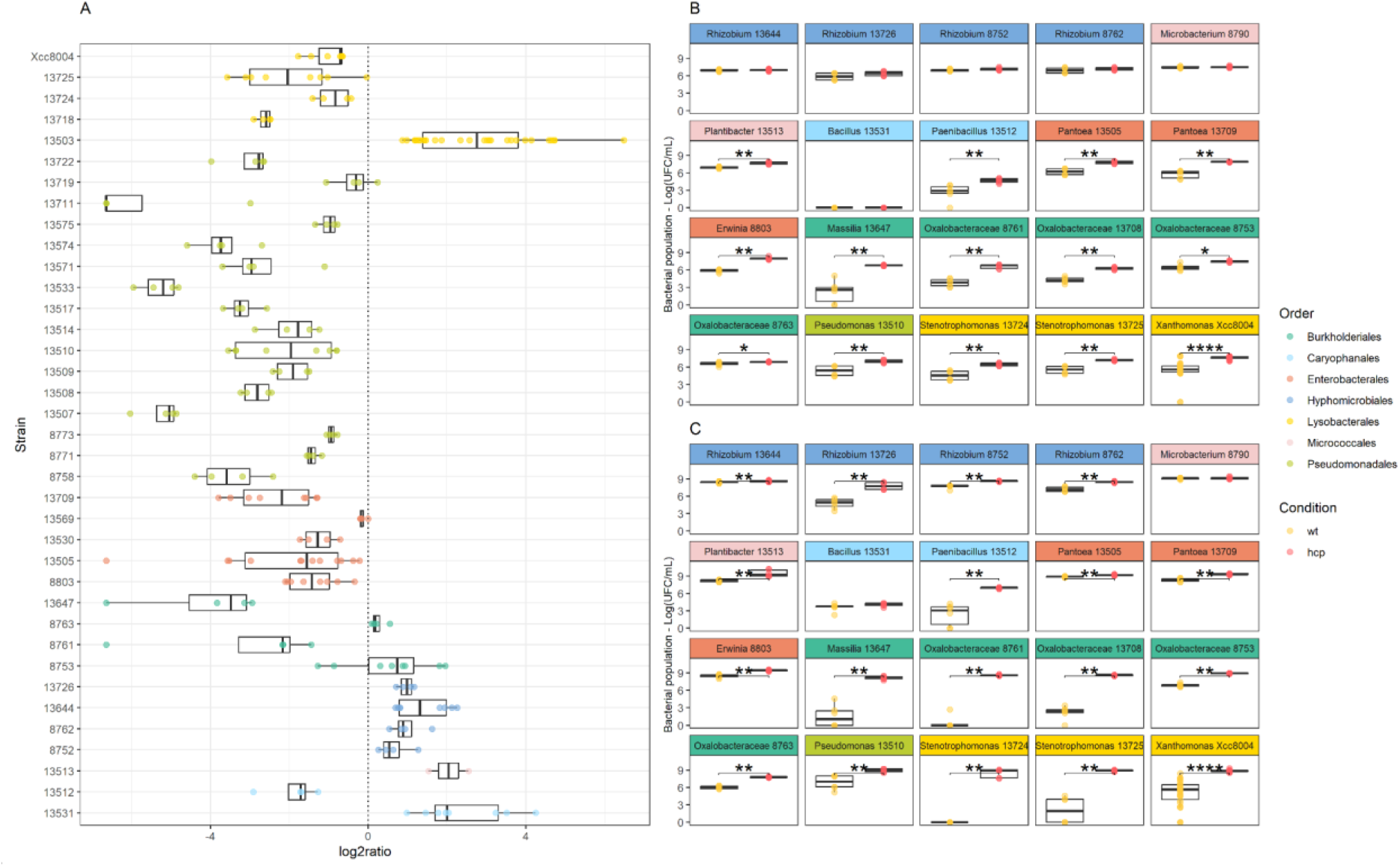
Strain sensitivity to T6SS-mediated competition of *S. rhizophila* CFBP13503. The strain relative abundance (log2ratio) between wild type vs *hcp* mutant confrontation in SynComs. The more negative value of log2ratio corresponds to a higher sensitivity to T6SS (**A**). Bacterial populations (CFU.ml^-1^) of some rifampicin-resistant strains were monitored after pairwise confrontation with *S. rhizophila* CFBP13503 wild-type (WT) or its isogenic T6SS-deficient mutant (*Δhcp*) in TSA10 medium for 6h (**B**) and 24h (**C**). Colony-forming units (CFU) were quantified on TSA10 supplemented with rifampicin. Six or 15 (Xcc8004) replicates are plotted. Statistical analyses were performed using Wilcoxon-Mann-Whitney Test (* p-value < 0.05; ** p-value <0.01; *** p-value < 0.001; **** p-value < 0.0001).

By comparing relative abundance of SynComs members at 6h and their cophenetic distances with CFBP13503 we detected a significant correlation (R^2^ = 0.25, *P* = 0.002, **Fig. 3B**). This relationship could be explained by a greater metabolic overlap for strains closer to *S. rhizophila* CFBP13503. Indeed, an inverse correlation (R^2^ = 0.26, *P* = 0.001) between changes in relative abundance of SynCom members and resource overlap with CFBP13503 was observed at 6h (**Fig. 3C**). At 24h, none of these correlations were significant (**Fig. S2**).

### The T6SS of *S. rhizophila* CFBP13503 inhibits the growth of a wide range of bacteria

Pairwise confrontation assays were performed between rifampicin-resistant variants of 20 selected SynCom members and either the WT or T6SS-deficient (Δ*hcp*) *S. rhizophila* CFBP13503 strains. These assays were designed to determine whether the observed T6SS-dependent shifts in relative abundance could be attributed to the sensitivity or resistance of the bacterial taxa to the T6SS of *S. rhizophila* CFBP13503. The data obtained with these confrontation assays at 6h were consistent with the estimated relative abundance for 17 of the 20 strains tested (**Fig. 4A** and **Fig. 4B, Table S1**). The three strains with conflicting data were all enriched in relative abundance within SynComs but were sensitive to the T6SS of CFBP13503 according to CFU count in pairwise confrontations assays. These strains belong to *Plantibacter* (CFBP13513) and *Oxalobacteraceae* (CFBP8753 and CFBP8763). Altogether these results indicate that the relative abundance measured at 6h with *gyrB* amplicon sequencing is a good first approximation for determining whether strains are sensitive or resistant to the T6SS of CFBP13503.

At 6h, the number of CFU of 14 strains was significantly (*P* <0.05) reduced by the T6SS of CFBP13503 (**Fig. 4B**). At 24 h, the growth of almost all tested strains (18 out of 20) were significantly (*P* <0.05) inhibited (**Fig. 4C**). Indeed, 4 strains belonging to *Hyphomicrobiales* order resistant at 6h showed population reduction after 24h confrontation with CFPB13503 WT strain.

Although there is a relationship between phylogenetic distance and the intensity of bacterial response to T6SS in CFBP13503, this is only partial and does not apply to all the bacteria targeted in this study. For instance, differences in response amplitude were observed within *Burkholderiales*, where 3 strains (CFBP13647, CFBP8761 and CFBP13708) were highly altered by the T6SS from *S. rhizophila* CFBP13503, whereas populations of CFBP8753 and CFBP8763 were less impacted even after 24 h of confrontation (**Fig. 4B**). This result demonstrates differences in bacterial sensitivity independently of the phylogenetic distance between competitor and predator. To rule out other potential factors, growth curve analysis in TSB10 medium was performed, revealing that the strains exhibited different growth rates. However, no correlation was observed between growth rate and T6SS sensitivity (**Fig. S3**), indicating that growth rate is not a determining factor for sensitivity.

Under our *in vitro* conditions, the T6SS of *S. rhizophila* CFBP13503 does not provide a competitive advantage, as the population sizes of the wild type (WT) and Δ*hcp* mutant did not differ significantly in pairwise confrontations at either of the two time points analyzed, except in interactions with *P. agglomerans* or other *Stenotrophomonas* strains. In these cases, the WT strain exhibited a significantly higher population size compared to the Δ*hcp* mutant (**Fig. S4**).

### *S. rhizophila* CFBP13503 T6SS influences specific bacterial taxa transmission from seed to seedling

The broad target range of the T6SS of *S. rhizophila* CFBP13503 observed *in vitro* questions (i) the role of this system in the transmission of this strain to seedling and (ii) its impact on the assembly of bacterial communities when seed and soil microbiota coalesce.

Since *S. rhizophila* CFBP13503 is efficiently transmitted to seedlings of radish under gnotobiotic conditions (33), we used this plant species to investigate its T6SS impact on microbiota transmission in living potting soil. We first inoculated seeds with CFBP13503 strains (WT or Δ*hcp*) and collected germinated seeds and seedlings. Under these experimental conditions, we confirmed the efficient transmission of *S. rhizophila* CFBP13503 on germinated seeds (average of 1.6×10^6^ cells) and seedlings (aerial: 2.4×10^5^ cells, root: 2.2×10^4^ cells, **Fig. 5A**). According to *gyrB* amplicon sequencing, CFBP13503 reached 15% (germinated seeds), 11% (seedling - aerial compartment) and 2% (seedling - root compartment) of the relative abundance of bacterial communities (**Fig. 5B**). No significant differences in CFBP13503 population size and relative abundance between WT and Δ*hcp* were detected in all studied habitats (**Fig. 5**), which suggest that the T6SS of this strain is not involved in seed-to-seedling transmission. It is worth noting that the differences in significance trends between CFU counts (Fig. 5A) and *gyrB* relative abundance (**Fig. 5B**) likely stem from the distinct nature of these measurements. While CFUs provide a direct, quantitative measure of viable bacterial cells, *gyrB* relative abundance represents the proportion of specific taxa within the total bacterial community, which can be influenced by shifts in overall community composition. Additionally, CFU counts reflect only cultivable cells, whereas *gyrB*-based metabarcoding also includes DNA from non-cultivable bacteria, potentially explaining discrepancies between the two approaches.

**Figure 5:**
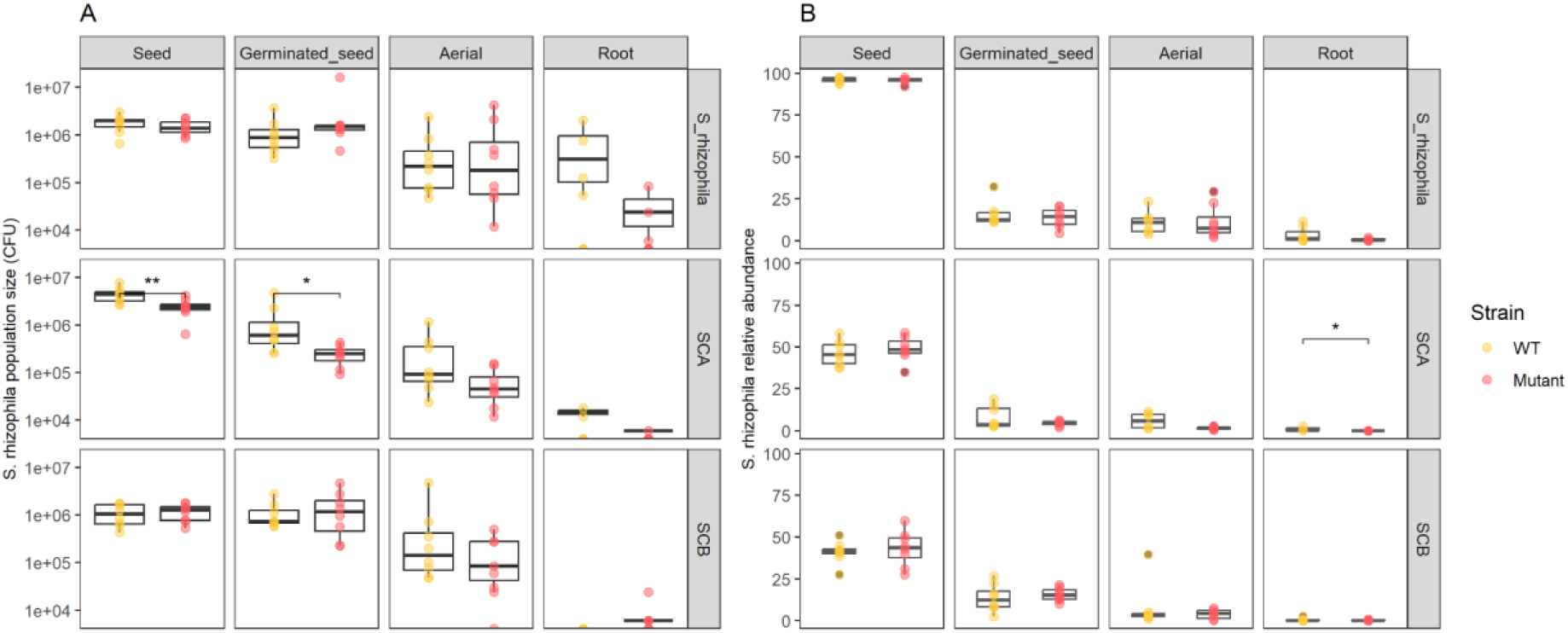
Abundance of *S. rhizophila* CFBP13503 during seed to seedling transmission. *S. rhizophila* CFBP13503 populations were enumerated on selective media (TSA10+Spe^50^+Amp^100^) media (**A**). The colors represent the inoculated *S. rhizophila* CFBP13503 strain WT (yellow) and Δ*hcp* (red). Data are plotted as CFU / individual. Relative abundance of *S. rhizophila* CFBP13503 according to *gyrB* amplicon sequencing (**B**). Statistical analyses were performed using Wilcoxon-Mann-Whitney test (* p-value < 0.05).

The high inoculation of CFBP13503 populations (10^6^ CFU per seed) within the native seed communities, generally around 10^3^ per seed (55), could explain why no competitive advantage for the wild-type strain was detected. A second experiment was therefore carried out through seed-inoculation of CFBP13503 strains (WT or Δ*hcp*) into two distinct synthetic communities (SCA and SCB). These two communities differ not only in Faith’s phylogenetic diversity (SCA: 1.9; SCB: 2.8) but also in their composition of T6SS-susceptible and -resistant strains, with SCA made up solely of susceptible strains and SCB made up of two susceptible strains and two resistant strains (**Table S2**). When CFBP13503 strains were seed-inoculated with SCA (low-diversity community composed of T6SS susceptible strains), CFBP13503 T6SS-deficient population sizes were significantly less abundant in seed and germinated-seeds compared to the WT population sizes (**Fig. 5A**). In contrast, for SCB (a higher-diversity community composed of two susceptible and two resistant strains), no significant differences were observed. By analyzing the relative abundance of SynComs members through *gyrB* amplicon sequencing, 4 strains showed significant (*P* < 0.05) increases in relative abundance at the germinated seed stage when coinoculated with the CFBP13503 T6SS-deficient mutant, compared to coinoculation with the wild-type strain (**Fig. 6**). Two of these strains belong to SCA, while the other two from SCB are two strains previously identified as T6SS-sensitive. These results indicate that some of the strains (4 out of 6) sensitive to the T6SS of *S. rhizophila* CFBP13503 *in vitro* were also sensitive to this secretion system *in planta*. However, this effect is fairly early, since the significant changes in relative abundance were all observed at the germinated seed stage.

**Figure 6.**
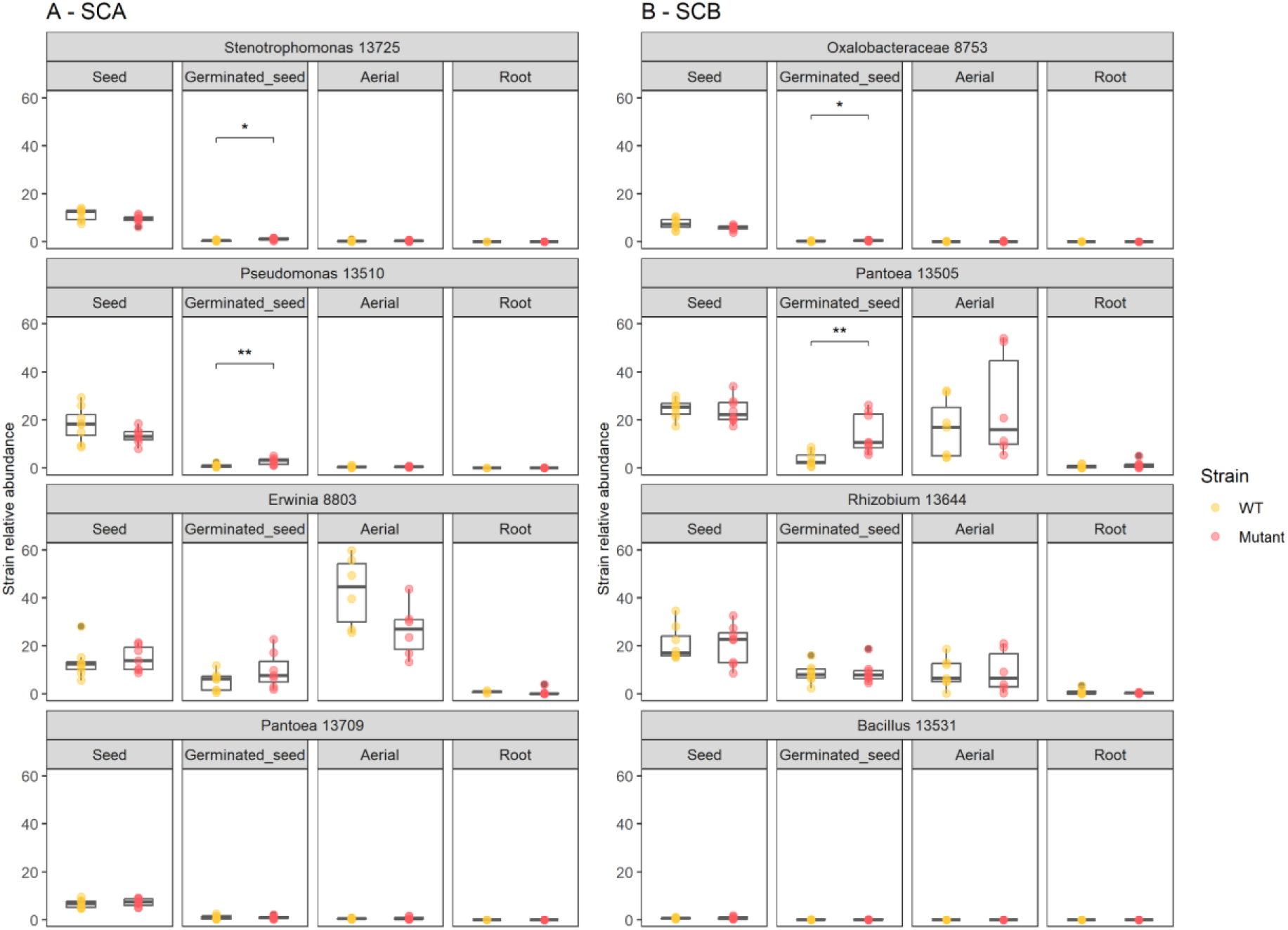
Relative abundance of SynCom members *in planta*. Relative abundance of seed-inoculated bacteria during interaction with *S. rhizophila* CFBP13503 strains (WT or Δ*hcp*). Statistical analyses were performed using Wilcoxon-Mann-Whitney test (* p-value < 0.05).

As SynCom members represented at best 50% of germinated-seed and seedling communities (**Fig. 6**), the impact of CFBP13503 T6SS on native members of the microbiota (i.e. non-inoculated, soil bacteria) was next investigated. No change in alpha-diversity (Shannon) and beta-diversity (weighted UniFrac distances) was detected. Relative abundance of each bacterial taxonomic rank (from phyla to ASV) was compared following inoculation with CFBP13503 WT strain or the T6SS-deficient mutant. We were able to detect very few taxa with significant changes in relative abundance (corrected p-value < 0.05). Only ASVs associated with *Bradyrhizobiaceae* had an increased relative abundance in the presence of the T6SS deficient mutant of CFBP13503 in the root compartment (**Fig. S6**).

## DISCUSSION

### T6SS is highly prevalent among seed-associated bacteria

T6SS is frequently found in Gram-negative bacteria with 17,183 putative T6SS gene clusters detected in 7,617 genomic sequences out of 26,573 as of November 2021 (44). Analysis of a diversity of bacterial strains isolated from radish, rapeseed and bean seed samples showed that 60% of genomic sequences had a T6SS secretion system predominantly affiliated to the T6SS^i^. This percentage rises to 75% when only Pseudomonodota are considered, a percentage similar to that previously described above for plant-associated Pseudomonodota (20).

### T6SS of *S. rhizophila* CFBP13503 targets a broad spectrum of bacterial species

The T6SS of *S. rhizophila* CFBP13503, which belongs to cluster i4B, is involved in growth inhibition of the plant pathogen *X. campestris* pv. *campestris* both *in vitro* and *in planta* (31). In the present work, we have demonstrated that this effect is involved in the *in vitro* inhibition of a broad spectrum of seed-borne bacteria belonging to *Enterobacterales*, *Lysobacterales*, *Burkholderiales* and *Pseudomonodales*. The diversity of competitors targeted by a single T6SS is not specific to *S. rhizophila* CFBP13503. For instance, the T6SS i4B of the plant-associated bacterial strains *Pseudomonas putida* KT2440 (56), *Burkholderia gladioli* NGJ1 (57) or *Paraburkholderia sabiae* LMG24235 (58) are involved in inhibition of a range of phytopathogenic agents including *Agrobacterium tumefaciens*, *Pectobacterium carotovorum*, *Pseudomonas syringae*, *Ralstonia solanacearum* and *Xanthomonas campestris*.

### The phylogenetic proximity of bacterial competitors to *S. rhizophila* CFBP13503 partly explains their susceptibility to its T6SS

Interference competition mediated through T6SS-contact dependent mechanisms was more prevalent among phylogenetically similar species. Moreover, according to predictions of resource acquisition pathways from bacterial strain genomes, bacteria that are phylogenetically and therefore also metabolically closer to *S. rhizophila* CFBP13503 are more likely to be susceptible to its T6SS. These relationships were observed only during the 6-hour confrontations. In multispecies biofilms, the confrontation of *P. putida* KT2440 with soil communities revealed similar results, where metabolic similarities led to an enhancement of T6SS antimicrobial activity (59). Taken together, these observations indicate that resource overlap drives only in part the evolution of T6SS-mediated interbacterial competition in a manner analogous to that observed for contact-independent inhibition (30).

### Immunity-dependent versus-independent mechanisms

The inhibition of a wide range of targeted competitor species by *S. rhizophila* T6SS could be explained by its large predicted T6SS effector repertoire (n=9), which cover a large variety of potential activities including phospholipases, amidases, pore-forming effectors and DNases (31). The concomitant delivery of a cocktail of effectors can provide a bet-hedging strategy to promote bacterial competitiveness in the face of unpredictable and variable environmental (60) like during transmission from seed to seedling. It is interesting to understand why certain bacterial orders are sensitive while others are resistant. It is highly probable that the resistance of strains to the T6SS of CFBP13503 is related to immunity-independent mechanisms (28). For instance, several monoderm strains tested in this work are resistant to the T6SS of CFBP13503. This resistance could be explained by the thick peptidoglycan layer of monoderms, which requires specific T6SS effectors to be perforated (61, 62) or the induction of survival mechanisms such as sporulation, which has been observed in *B. subtilis* in response to the secretion of a muramidase-type effector secreted by *P. chlororaphis* (63). Concerning *Burkholderiales*, while certain strains exhibit resistance within the communities, they paradoxically show susceptibility when individually confronted with *S. rhizophila* CFBP13503. The varied responses of *Burkholderiales* could be attributed to collective protection within communities. This type of protection has been observed in *E. coli*, where the secretion of exopolysaccharides (EPS) serves to shield unproductive EPS strains from the T6SS of *Acinetobacter baylyi* (64). Another explanation could be the existence of immunity genes in some targeted bacteria that protect from some toxical T6SS effectors (T6Es) of *S. rhizophila* CFBP1350. Such hypotheses will be investigated by looking for putative interactions between T6Es and immunity proteins from surrounding bacteria.

### SynComs response to T6SS in CFBP13503 not explained by T6SS richness

T6SS can be assembled in different ways from random firing (e.g. *Vibrio cholerae* (65)) to specific response to T6SS attack (e.g. *Pseudomonas aeruginosa* (66)). The T6SS i4B of *S. rhizophila* CFBP13503 possesses an ortholog of TslA (31), a periplasmic protein involved in T6SS assembly at the site of contact between neighboring cells (67). Hence it is likely that the T6SS of *S. rhizophila* CFBP13503 is activated during contact with competitor cells in our *in vitro* confrontation assays. By defining synthetic communities with different T6SS diversity, we anticipated a response from T6SS-positive strains that have notably a tit-for-tat strategy (i.e. some *Pseudomonas* strains with a T6SS i3). This response should ultimately limit the competitive advantage of CFBP13503 in these T6SS-positive communities. According to *gyrB* amplicon sequencing, the richness of T6SS in SynComs was not correlated to CFBP13503 relative abundance. Several factors may explain this observation, including the high ratio of CFBP13503 in relation to the competitors, a large spectrum of immunity against other bacteria T6SS effectors or the T6SS systems of the other bacterial competitors might not be actively functioning under *in vitro* conditions. Indeed, the T6SS is a tightly regulated mechanism, with its expression being responsive to an array of biotic and/or abiotic factors (68–70).

### T6SS impact on bacterial community assembly during seed to seedling transmission depends on the inoculated SynCom composition on seed

We investigated how *S. rhizophila* CFBP13503 T6SS impacts bacterial community dynamics during seed-to-seedling transmission, aiming to observe bacteria responses within their natural environment. The inoculation of bacterial SynComs is the most reliable way to homogenize seed colonization compared to the highly variable single seed microbiota (7, 55). Using *gyrB* metabarcoding, we tracked the transmission of two synthetic communities and soil-associated bacteria across various stages of plant development, encompassing seeds, germinated seeds, and seedlings. The different SynCom members successfully colonized the seeds, resulting in distinct community compositions at the seed stage. However, a convergence in community composition was detected upon seed germination, which reflects a preferential colonization of this habitat by soil-borne taxa as already observed in rapeseed and wheat (5, 8). While the T6SS of *S. rhizophila* CFBP13503 was not an important driver of whole community composition, changes in relative abundances of SynCom members in germinating seeds and some soil-borne taxa (i.e. *Bradyrhizobiaceae*) in roots were detected. The impact of T6SS on the relative abundance of susceptible inoculated strains resulted in an increase in the population and relative abundance of *S. rhizophila* CFBP13503, demonstrating that T6SS can provide a competitive advantage depending on the susceptibility of bacterial competitors. This competitive advantage enhances the bacterium ability to persist in the microbiota over time. A stable to increased persistence potentially maintains or increases the probability of outcompeting other T6SS-sensitive bacteria. This persistence could be crucial for establishing stable and beneficial interactions with the plant host. However, a subset of SynCom members (CFBP13709, CFBP8803, and CFBP8753) remained unaffected. This disparity in response could be attributed to the contrasting environments of *in vitro* confrontation assays, where direct cell-to-cell contact occurs within a confined space, and the more complex spatial dynamics introduced by seed germination and plant growth. The intricate nature of seed germination and plant growth introduces substantial spatial disruptions, a phenomenon demonstrated to increasingly favor sensitive strains that can evade contact with T6SS-producing cells (71). Furthermore, to discern the direct impacts of T6SS on a bacterial member, co-localization is essential (72). Only specific plant habitats may be affected by the T6SS of competitors as it was demonstrated for the gallobiome of tomato induced by *Agrobacterium*, but only during summer time of gall development (73). The precise nature of the impact of *S. rhizophila* T6SS on individual bacterial taxa, whether direct or indirect, remains to be determined. Nonetheless, our study elucidates a clear pattern: while T6SS exerts a negative and specific impact on certain bacterial taxa, it simultaneously creates a favorable environment for other taxa. These changes in bacterial taxa distribution have cascading effects on bacterial interactions, resulting in distinct community dynamics.

## ACKNOWLEDGEMENTS

This work was funded by the TypeSEEDS project (INRAE: Plant Health and Environment division, HoloFlux Metaprogram) and the Région des Pays de la Loire (RFI “Objectif Végétal”). We thank the CIRM-CFBP (https://doi.org/10.15454/E8XX-4Z18) for providing access to strains of their culture collection and Muriel Bahut (ANAN platform, SFR QuaSaV) for amplicon sequencing.

## Notes

### Competing Interest Statement

The authors have declared no competing interest.

### Summary of Updates

We (i) have more explicitly justified the choice and design of SynComs, (ii) add quantitative values (qPCR and colony-forming units) to amplicon analysis. With regard to the expression of T6SS, the observed phenotypes (e.g. change in community size and composition; modulation of bacterial population size in pairwise confrontation) indicate that the T6SS of S. rhizophila CFBP13503 is expressed under these experimental conditions. The mechanisms involved in regulating this system are currently being addressed in a subsequent study.

