## Supplemental for "T6SS-mediated competition by *Stenotrophomonas rhizophila* shapes seed-borne bacterial communities and seed-to-seedling transmission dynamics"

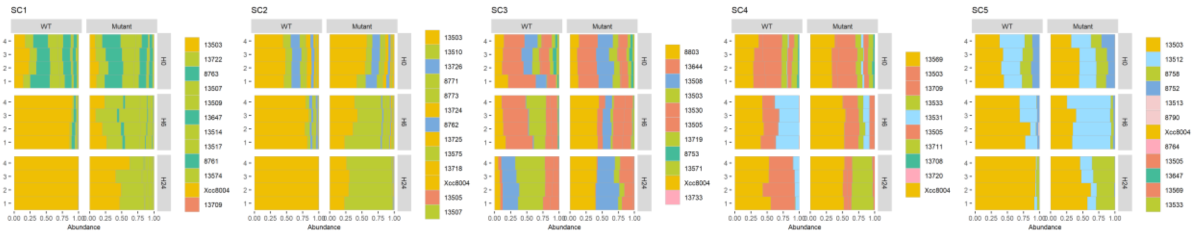


**Figure S1. T6SS impact on the structure of bacterial synthetic communities *in vitro*.**
Relative abundance of bacterial taxa within five different synthetic communities (SynCom) at different confrontation times (0h, 6h, and 24h) while competing with the wild-type strain of *S. rhizophila* CFBP13503 (WT) or the T6SS-deficient mutant Δ*hcp*.

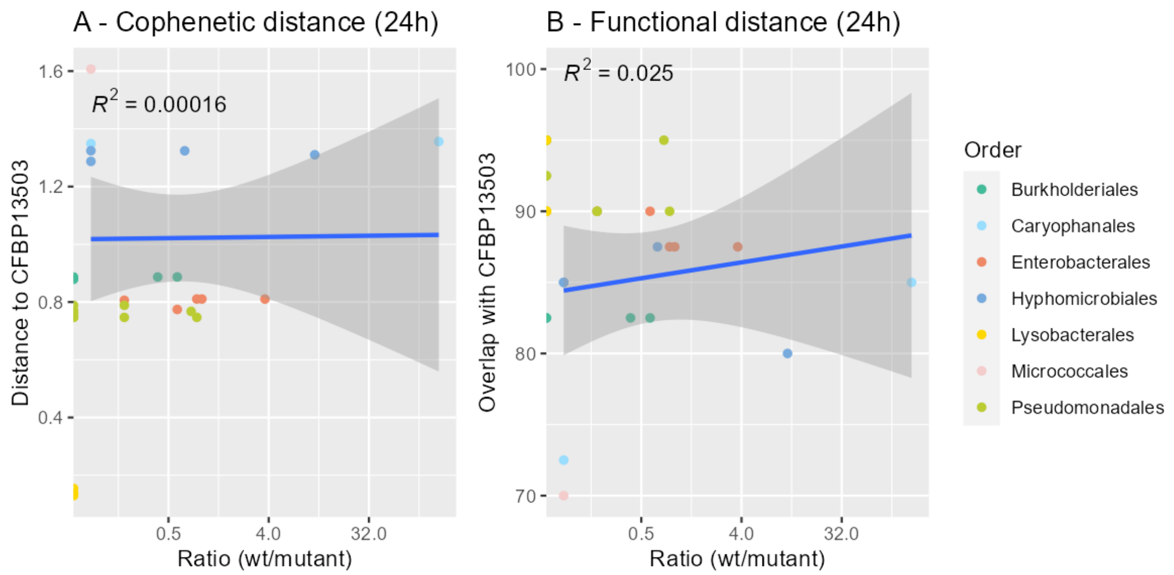


**Figure S2 Relationship between phylogenetic/functional distances and RA at 24h confrontation of SynCom members.** Correlation between changes in relative abundance of SynCom members at 24h and their cophenetic distances to CFBP13503 (**A**). Correlation between changes in relative abundance of SynCom members at 24h and their resource overlap with CFBP13503 (**B**).

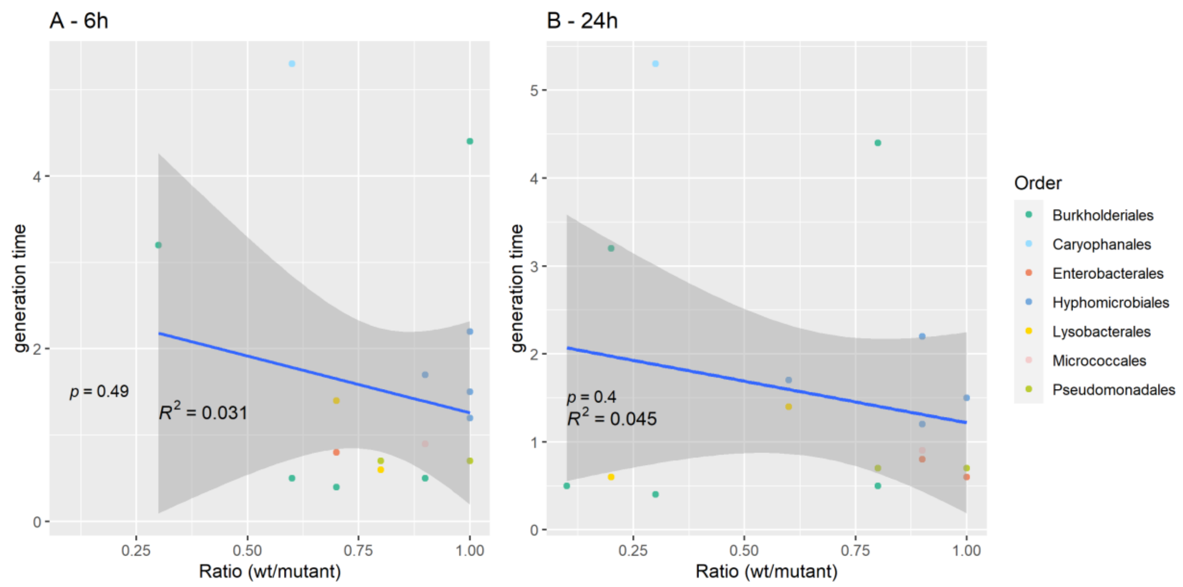


**Figure S3. Relationship between strain sensitivity to T6SS and strain growth rate at 6h and 24h of confrontation with CFP13503 (WT) and Δ*hcp*.**

Strain sensitivity is reported as the ratio (LogCFU WT/Δ*hcp*) and growth rate as generation time (h) in TSB1/10.

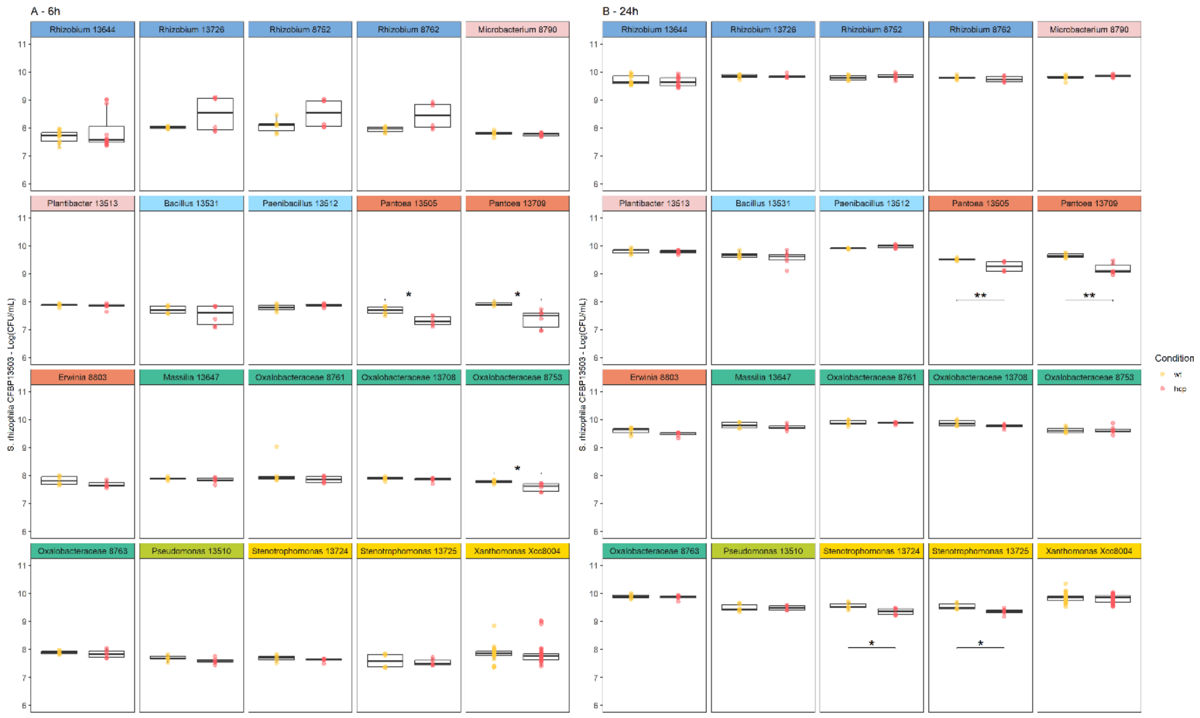


**Figure S4. Population dynamics of *S. rhizophila* CFBP13503 and T6SS-deficient mutant *Δhcp* during *in vitro* confrontation with seed-borne bacterial strains**. *S. rhizophila* populations (CFU.ml$^{-1}$) were monitored after confrontation with rifampicin-resistant strains in TSA10 medium for 6h (A) and 24h (B). Colony-forming units (CFU) were quantified on TSA10 supplemented with spectinomycin and ampicillin. Six replicates are plotted. Statistical analyses were performed using Wilcoxon-Mann-Whitney Test (* p-value < 0.05).

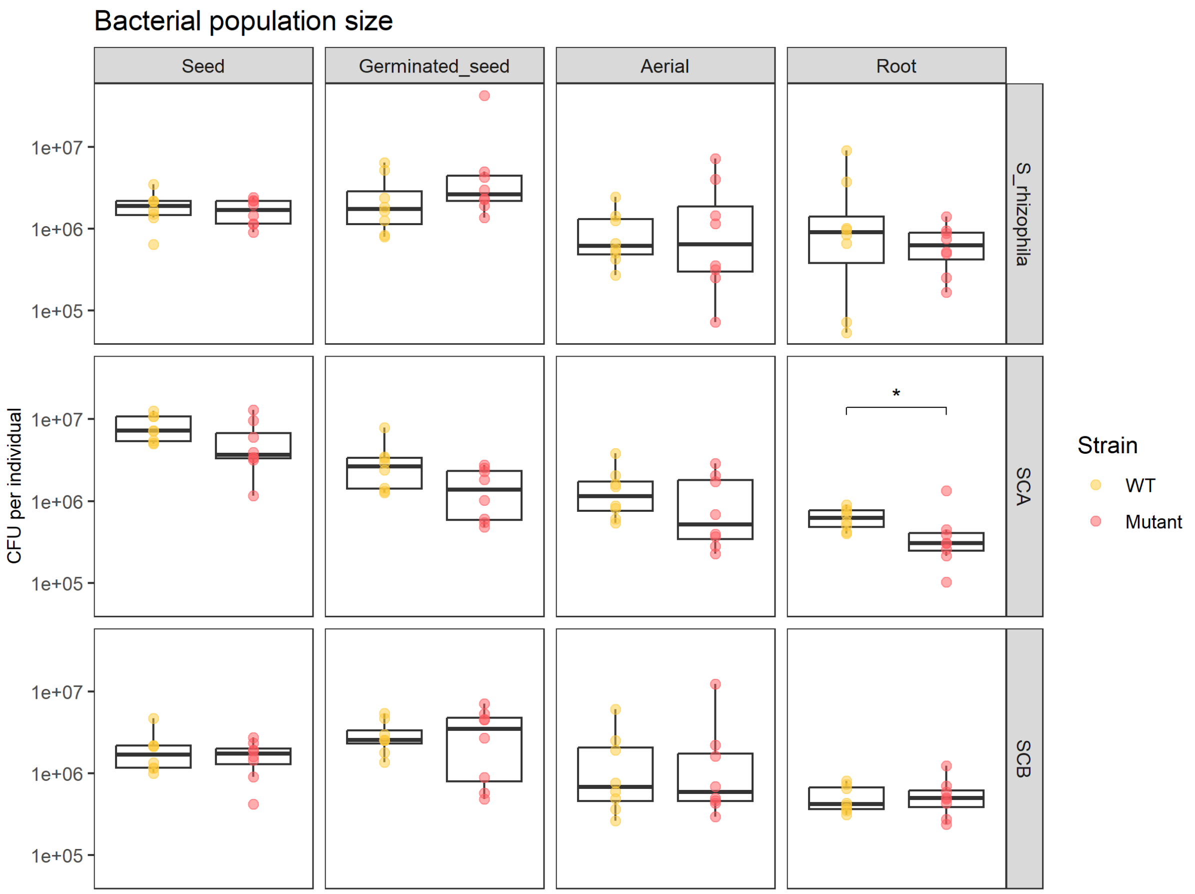


**Figure S5. Abundance of total bacteria during seed to seedling transmission**. Bacterial community size was enumerated on TSA10 medium. The colors represent the initial seed inoculation with *S. rhizophila* CFBP13503 strain WT (yellow) and Δ*hcp* (red). Data are plotted as CFU / individual. Statistical analyses were performed using Wilcoxon-Mann-Whitney test (* p-value < 0.05).

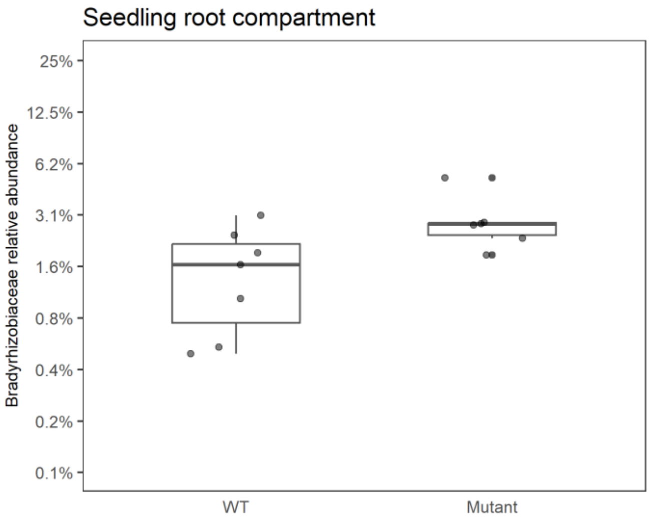


**Figure S6. Relative abundance of ASVs affiliated to *Bradyrhizobiaceae* in roots following seed-inoculations of *S. rhizophila* CFBP13503 strains (WT or T6SS-deficient mutant).** Relative abundance data were log2 transformed for display purposes.

**Table S1. Investigation of T6SS presence within genome sequences of seed and seedling-associated bacterial strains.**

| CFBP | Taxonomy | Biosample | Host | Compartmen | Year | Plot | Genome | GC% | CDSs | T6SS | i1 | i2 | i3 | i4a | i4b | i5 | iii | Bioproject |
|---|---|---|---|---|---|---|---|---|---|---|---|---|---|---|---|---|---|---|
| 7767 | Xanthomonas citri pv. fuscans | SAMN06829140 | Phaseolus vulgaris | Seed | 2009 | Cameroun | 5297402 | 65 | 4516 | 2 | | 1 | | | 1 | | | PRJNA383319 |
| 8751 | Frigoribacterium | SAMN16237779 | Brassica napus Tenor | Seed | 2017 | IGEPP | 3539903 | 71 | 3293 | 0 | | | | | | | | PRJNA665156 |
| 8752 | Rhizobium | SAMN16237344 | Brassica napus Boston | Seed | 2017 | IGEPP | 4888650 | 60 | 4580 | 0 | | | | | | | | PRJNA665156 |
| 8753 | Oxalobacteraceae | SAMN16237354 | Brassica napus Boston | Seed | 2017 | IGEPP | 5167451 | 64 | 4430 | 1 | | | | | 1 | | | PRJNA665156 |
| 8754 | Frigoribacterium | SAMN16237571 | Brassica napus Major | Seed | 2017 | IGEPP | 3385781 | 73 | 3080 | 0 | | | | | | | | PRJNA665156 |
| 8755 | Oxalobacteraceae | SAMN16237644 | Brassica napus Major | Seed | 2017 | IGEPP | 5220625 | 64 | 4513 | 1 | | | | | 1 | | | PRJNA665156 |
| 8756 | Pantoea agglomerans | SAMN16237653 | Brassica napus Major | Seed | 2017 | IGEPP | 5029763 | 55 | 4637 | 2 | | 1 | 1 | | | | | PRJNA665156 |
| 8757 | Aeromicrobium | SAMN16237654 | Brassica napus Major | Seed | 2017 | IGEPP | 3757853 | 71 | 3670 | 0 | | | | | | | | PRJNA665156 |
| 8758 | Pseudomonas putida group | SAMN16237690 | Brassica napus Major | Seed | 2017 | IGEPP | 4663592 | 62 | 4189 | 0 | | | | | | | | PRJNA665156 |
| 8759 | Frigoribacterium | SAMN16237713 | Brassica napus Mohican | Seed | 2017 | IGEPP | 3401111 | 71 | 3141 | 0 | | | | | | | | PRJNA665156 |
| 8760 | Sphingomonas | SAMN16237746 | Brassica napus Mohican | Seed | 2017 | IGEPP | 4302246 | 67 | 3972 | 1 | | | | | | 1 | | PRJNA665156 |
| 8761 | Oxalobacteraceae | SAMN16237762 | Brassica napus Tenor | Seed | 2017 | IGEPP | 5211781 | 64 | 4452 | 1 | | | | | 1 | | | PRJNA665156 |
| 8762 | Rhizobium | SAMN16237769 | Brassica napus Tenor | Seed | 2017 | IGEPP | 4436340 | 57 | 4279 | 0 | | | | | | | | PRJNA665156 |
| 8763 | Oxalobacteraceae | SAMN16237778 | Brassica napus Tenor | Seed | 2017 | IGEPP | 5498755 | 63 | 4693 | 2 | | | | | 2 | | | PRJNA665156 |
| 8764 | Sphingomonas | SAMN16237800 | Brassica napus Tenor | Seed | 2017 | IGEPP | 4345566 | 66 | 3880 | 0 | | | | | | | | PRJNA665156 |
| 8765 | Sphingomonas | SAMN16237811 | Brassica napus Zorro | Seed | 2017 | IGEPP | 4387101 | 66 | 3941 | 0 | | | | | | | | PRJNA665156 |
| 8766 | Frigoribacterium | SAMN16237812 | Brassica napus Zorro | Seed | 2017 | IGEPP | 3559033 | 72 | 3254 | 0 | | | | | | | | PRJNA665156 |
| 8767 | Erw inia persicina | SAMN16237479 | Brassica napus Express | Seed | 2017 | IGEPP | 5182457 | 55 | 4829 | 3 | | 3 | | | | | | PRJNA665156 |
| 8768 | Pseudomonas fluorescens subgroup | SAMN16237480 | Brassica napus Express | Seed | 2017 | IGEPP | 6155614 | 60 | 5556 | 2 | 1 | | 1 | | | | | PRJNA665156 |
| 8769 | Pseudomonas putida group | SAMN16237574 | Brassica napus Major | Seed | 2017 | IGEPP | 5709324 | 60 | 5245 | 0 | | | | | | | | PRJNA665156 |
| 8770 | Pseudomonas putida group | SAMN16237655 | Brassica napus Major | Seed | 2017 | IGEPP | 4654033 | 62 | 4125 | 0 | | | | | | | | PRJNA665156 |
| 8771 | Pseudomonas putida group | SAMN16237697 | Brassica napus Major | Seed | 2017 | IGEPP | 4765370 | 62 | 4232 | 1 | 1 | | | | | | | PRJNA665156 |
| 8772 | Pseudomonas | SAMN16237710 | Brassica napus Major | Seed | 2017 | IGEPP | 5879355 | 61 | 5162 | 3 | 1 | 1 | | | 1 | | | PRJNA665156 |
| 8773 | Pseudomonas putida group | SAMN16237754 | Brassica napus Mohican | Seed | 2017 | IGEPP | 4654405 | 62 | 4121 | 0 | | | | | | | | PRJNA665156 |
| 8774 | Pantoea agglomerans | SAMN16237802 | Brassica napus Zorro | Seed | 2017 | IGEPP | 5464831 | 55 | 5207 | 2 | | 1 | 1 | | | | | PRJNA665156 |
| 8775 | Plantibacter | SAMN16237042 | Phaseolus vulgaris Flavert | Flow er | 2016 | FNAMS – 49 | 4397458 | 69 | 4069 | 0 | | | | | | | | PRJNA665156 |
| 8776 | Pseudomonas fluorescens subgroup | SAMN16237045 | Phaseolus vulgaris Flavert | Flow er | 2016 | FNAMS – 49 | 6046465 | 60 | 5454 | 3 | 1 | 1 | 1 | | | | | PRJNA665156 |
| 8777 | Pseudomonas syringae group | SAMN16237050 | Phaseolus vulgaris Flavert | Flow er | 2016 | FNAMS – 49 | 6139145 | 62 | 5479 | 2 | 1 | | 1 | | | | | PRJNA665156 |
| 8778 | Pseudomonas fluorescens subgroup | SAMN16237054 | Phaseolus vulgaris Flavert | Pods | 2016 | FNAMS – 49 | 6013069 | 60 | 5290 | 3 | 1 | 1 | 1 | | | | | PRJNA665156 |
| 8779 | Pseudomonas syringae group | SAMN16237056 | Phaseolus vulgaris Flavert | Pods | 2016 | FNAMS – 49 | 6140205 | 62 | 5475 | 2 | 1 | | 1 | | | | | PRJNA665156 |
| 8780 | Pseudomonas orientalis | SAMN16237064 | Phaseolus vulgaris Flavert | Seeds | 2016 | FNAMS – 49 | 5804252 | 60 | 5109 | 2 | 1 | | 1 | | | | | PRJNA665156 |
| 8781 | Pseudomonas orientalis | SAMN16237067 | Phaseolus vulgaris Flavert | Seeds | 2016 | FNAMS – 49 | 5802148 | 60 | 5109 | 2 | 1 | | 1 | | | | | PRJNA665156 |
| 8782 | Xanthomonas campestris pv. campestris | SAMN16237897 | Raphanus sativus Flamboyant5 | Seed | 2014 | FNAMS – 49 | 5076693 | 65 | 4257 | 0 | | | | | | | | PRJNA665156 |
| 8783 | Pantoea agglomerans | SAMN16237909 | Raphanus sativus Flamboyant5 | Flow er | 2016 | FNAMS – 49 | 5124033 | 55 | 4840 | 2 | | 1 | 1 | | | | | PRJNA665156 |
| 8784 | Pantoea agglomerans | SAMN16237910 | Raphanus sativus Flamboyant5 | Flow er | 2016 | FNAMS – 49 | 4755237 | 55 | 4336 | 2 | | 1 | 1 | | | | | PRJNA665156 |
| 8785 | Pantoea agglomerans | SAMN16237912 | Raphanus sativus Flamboyant5 | Flow er | 2016 | FNAMS – 49 | 5044125 | 55 | 4709 | 2 | | 1 | 1 | | | | | PRJNA665156 |
| 8786 | Pantoea agglomerans | SAMN16237914 | Raphanus sativus Flamboyant5 | Seed | 2016 | FNAMS – 49 | 4928760 | 55 | 4576 | 1 | | | 1 | | | | | PRJNA665156 |
| 8787 | Pseudomonas fluorescens subgroup | SAMN16237931 | Raphanus sativus Flamboyant5 | Seed | 2016 | FNAMS – 49 | 6016319 | 60 | 5295 | 3 | 1 | 1 | 1 | | | | | PRJNA665156 |
| 8788 | Pseudomonas fluorescens subgroup | SAMN16237973 | Raphanus sativus Flamboyant5 | Seed | 2016 | FNAMS – 49 | 6045815 | 60 | 5451 | 3 | 1 | 1 | 1 | | | | | PRJNA665156 |
| 8789 | Pseudomonas fluorescens subgroup | SAMN16238009 | Raphanus sativus Flamboyant5 | Seed | 2016 | FNAMS – 49 | 6046475 | 60 | 5453 | 3 | 1 | 1 | 1 | | | | | PRJNA665156 |
| 8790 | Pseudomonas fluorescens subgroup | SAMN16238272 | Raphanus sativus Flamboyant5 | Seed | 2016 | FNAMS – 49 | 3445937 | 70 | 3168 | 0 | | | | | | | | PRJNA665156 |
| 8791 | Pantoea agglomerans | SAMN16238273 | Raphanus sativus Flamboyant5 | Seed | 2016 | FNAMS – 49 | 4905192 | 55 | 4552 | 2 | | 1 | 1 | | | | | PRJNA665156 |
| 8792 | Pantoea agglomerans | SAMN16238279 | Raphanus sativus Flamboyant5 | Seed | 2016 | FNAMS – 49 | 5239108 | 55 | 4959 | 2 | | 1 | 1 | | | | | PRJNA665156 |
| 8793 | Pseudomonas orientalis | SAMN16238286 | Raphanus sativus Flamboyant5 | Seed | 2016 | FNAMS – 49 | 5799590 | 60 | 5105 | 2 | 1 | | 1 | | | | | PRJNA665156 |
| 8794 | Microbacterium | SAMN16238307 | Raphanus sativus Flamboyant5 | Seed | 2016 | FNAMS – 49 | 3502222 | 70 | 3236 | 0 | | | | | | | | PRJNA665156 |
| 8795 | Erw inia persicina | SAMN16238309 | Raphanus sativus Flamboyant5 | Seed | 2016 | FNAMS – 49 | 5148964 | 55 | 4784 | 1 | | 1 | | | | | | PRJNA665156 |
| 8796 | Pseudomonas orientalis | SAMN16238321 | Raphanus sativus Flamboyant5 | Seed | 2016 | FNAMS – 49 | 5802756 | 60 | 5101 | 2 | 1 | | 1 | | | | | PRJNA665156 |
| 8797 | Erw inia persicina | SAMN16238323 | Raphanus sativus Flamboyant5 | Seed | 2016 | FNAMS – 49 | 4806053 | 56 | 4361 | 2 | | 2 | | | | | | PRJNA665156 |
| 8798 | Plantibacter | SAMN16238397 | Raphanus sativus Flamboyant5 | Seed | 2016 | FNAMS – 49 | 4301372 | 69 | 3949 | 0 | | | | | | | | PRJNA665156 |
| 8799 | Pseudomonas viridiflava | SAMN16237943 | Raphanus sativus Flamboyant5 | Seed | 2016 | FNAMS – 49 | 6035893 | 59 | 5388 | 2 | 1 | | 1 | | | | | PRJNA665156 |
| 8800 | Pantoea agglomerans | SAMN16238278 | Raphanus sativus Flamboyant5 | Seed | 2016 | FNAMS – 49 | 5117730 | 55 | 4806 | 2 | 1 | 1 | | | | | | PRJNA665156 |
| 8801 | Microbacterium | SAMN16237052 | Phaseolus vulgaris Flavert | Flow er | 2016 | FNAMS – 49 | 3442739 | 70 | 3159 | 0 | | | | | | | | PRJNA665156 |
| 8802 | Pseudomonas viridiflava | SAMN16238274 | Raphanus sativus Flamboyant5 | Seed | 2016 | FNAMS – 49 | 5947296 | 59 | 5265 | 2 | 1 | | 1 | | | | | PRJNA665156 |
| 8803 | Erw inia persicina | SAMN16237911 | Raphanus sativus Flamboyant5 | Flow er | 2016 | FNAMS – 49 | 4897238 | 56 | 4491 | 2 | | 2 | | | | | | PRJNA665156 |
| 8804 | Plantibacter | SAMN16238287 | Raphanus sativus Flamboyant5 | Seed | 2016 | FNAMS – 49 | 4227001 | 69 | 3949 | 0 | | | | | | | | PRJNA665156 |
| 8977 | Pseudomonas colepterorum | SAMN38288732 | Phaseolus vulgaris Linex | Seedling | 2021 | FNAMS – 32 | 4820021 | 62 | 4264 | 0 | | | | | | | | PRJNA1041598 |
| 8978 | Lelliottia | SAMN38288733 | Phaseolus vulgaris Vanilla | Seedling | 2021 | FNAMS – 32 | 4442944 | 54 | 4027 | 3 | | 2 | 1 | | | | | PRJNA1041598 |
| 8979 | Pseudomonas syringae | SAMN38288734 | Phaseolus vulgaris Vanilla | Seedling | 2021 | FNAMS – 49 | 6056903 | 59 | 5146 | 1 | | | | | 1 | | | PRJNA1041598 |
| 8980 | Stenotrophomonas | SAMN38288735 | Phaseolus vulgaris Caprice | Seedling | 2021 | FNAMS – 32 | 3975215 | 66 | 3443 | 0 | | | | | | | | PRJNA1041598 |
| 8982 | Pseudomonas colepterorum | SAMN38288736 | Phaseolus vulgaris Caprice | Seed | 2021 | FNAMS – 32 | 4754523 | 62 | 4164 | 0 | | | | | | | | PRJNA1041598 |
| 8983 | Pseudomonas viridiflava | SAMN38288737 | Phaseolus vulgaris Deezer | Seed | 2021 | FNAMS – 49 | 5866520 | 59 | 5188 | 2 | 1 | | 1 | | | | | PRJNA1041598 |
| 8984 | Pseudomonas rhizosphaerae | SAMN38288738 | Phaseolus vulgaris Deezer | Seedling | 2021 | FNAMS – 49 | 4572494 | 62 | 4073 | 0 | | | | | | | | PRJNA1041598 |
| 8985 | Pseudomonas viridiflava | SAMN38288739 | Phaseolus vulgaris Vanilla | Seed | 2021 | FNAMS – 49 | 5852582 | 59 | 5155 | 1 | | | | | 1 | | | PRJNA1041598 |
| 8986 | Kosakonia | SAMN38288740 | Phaseolus vulgaris Linex | Seed | 2021 | FNAMS – 49 | 4808856 | 56 | 4393 | 1 | | | 1 | | | | | PRJNA1041598 |
| 8987 | Leclercia | SAMN38288741 | Phaseolus vulgaris Vanilla | Seedling | 2021 | FNAMS – 32 | 4836644 | 56 | 4496 | 1 | | 1 | | | | | | PRJNA1041598 |
| 8988 | Pantoea agglomerans | SAMN38288742 | Phaseolus vulgaris Deezer | Seedling | 2021 | FNAMS – 32 | 4810088 | 55 | 4403 | 2 | | 1 | 1 | | | | | PRJNA1041598 |
| 8989 | Pantoea agglomerans | SAMN38288743 | Phaseolus vulgaris Vanilla | Seedling | 2021 | FNAMS – 49 | 4766298 | 55 | 4375 | 2 | | 1 | 1 | | | | | PRJNA1041598 |
| 8990 | Siccibacter turicensis | SAMN38288744 | Phaseolus vulgaris Vanilla | Seedling | 2021 | FNAMS – 32 | 4131421 | 58 | 3760 | 3 | 1 | 1 | 1 | | | | | PRJNA1041598 |
| 8991 | Pseudomonas viridiflava | SAMN38288745 | Phaseolus vulgaris Linex | Seedling | 2021 | FNAMS – 32 | 5864384 | 59 | 5187 | 2 | 1 | | 1 | | | | | PRJNA1041598 |
| 8992 | Pseudomonas fluorescens subgroup | SAMN38288746 | Phaseolus vulgaris Caprice | Seedling | 2021 | FNAMS – 32 | 6313504 | 60 | 5708 | 3 | 1 | 1 | 1 | | | | | PRJNA1041598 |
| 8993 | Sphingomonas | SAMN38288747 | Phaseolus vulgaris Linex | Seedling | 2021 | FNAMS – 49 | 3824895 | 67 | 3436 | 0 | | | | | | | | PRJNA1041598 |
| 8994 | Stenotrophomonas | SAMN38288748 | Phaseolus vulgaris Caprice | Seedling | 2021 | FNAMS – 32 | 4250922 | 67 | 3720 | 0 | | | | | | | | PRJNA1041598 |
| 8995 | Enterobacter | SAMN38288749 | Phaseolus vulgaris Linex | Seedling | 2021 | FNAMS – 49 | 5194180 | 55 | 4735 | 1 | | 1 | | | | | | PRJNA1041598 |
| 8996 | Chryseobacterium | SAMN38288750 | Phaseolus vulgaris Vanilla | Seedling | 2021 | FNAMS – 49 | 4439189 | 34 | 4011 | 1 | | | | | | | 1 | PRJNA1041598 |
| 9000 | Pantoea agglomerans | SAMN38288751 | Phaseolus vulgaris Flavert | Seedling | 2021 | FNAMS – 49 | 4725785 | 55 | 4300 | 2 | | 1 | 1 | | | | | PRJNA1041598 |
| 9003 | Pseudomonas atacamensis | SAMN38288752 | Phaseolus vulgaris Contender | Seedling | 2021 | FNAMS – 32 | 5859533 | 60 | 5218 | 1 | | 1 | | | | | | PRJNA1041598 |
| 9006 | Stenotrophomonas rhizophila | SAMN38288753 | Phaseolus vulgaris Facila | Seedling | 2021 | FNAMS – 32 | 4752770 | 67 | 4124 | 1 | | | | | 1 | | | PRJNA1041598 |
| 9009 | Bacillus | SAMN38288754 | Phaseolus vulgaris Flavert | Seedling | 2021 | FNAMS – 32 | 5684090 | 40 | 5591 | 0 | | | | | | | | PRJNA1041598 |
| 9010 | Priestia megaterium | SAMN38288755 | Phaseolus vulgaris Flavert | Seedling | 2021 | FNAMS – 32 | 5857104 | 37 | 6029 | 0 | | | | | | | | PRJNA1041598 |
| 9011 | Curtobacterium | SAMN38288756 | Phaseolus vulgaris Vezer | Seedling | 2021 | FNAMS – 49 | 3616265 | 71 | 3424 | 0 | | | | | | | | PRJNA1041598 |
| 9012 | Massilia | SAMN38288757 | Phaseolus vulgaris Flavert | Seedling | 2021 | FNAMS – 49 | 5989676 | 65 | 5153 | 2 | | | 1 | | 1 | | | PRJNA1041598 |
| 9014 | Bacillus thuringiensis | SAMN38288758 | Phaseolus vulgaris Flavert | Seedling | 2021 | FNAMS – 32 | 5836202 | 35 | 5769 | 0 | | | | | | | | PRJNA1041598 |
| 9016 | Pseudoclavibacter | NA | Phaseolus vulgaris Facila | Seedling | 2021 | FNAMS – 32 | 3800658 | 68 | 3544 | 0 | | | | | | | | #N/A |
| 9017 | Sphingomonas | NA | Phaseolus vulgaris Facila | Seedling | 2021 | FNAMS – 32 | 4316920 | 66 | 3804 | 0 | | | | | | | | #N/A |
| 9019 | Sphingomonas | SAMN38288759 | Phaseolus vulgaris Vezer | Seedling | 2021 | FNAMS – 32 | 4245651 | 66 | 3796 | 0 | | | | | | | | PRJNA1041598 |
| 9020 | Endobacterium cereale | SAMN38288760 | Phaseolus vulgaris Flavert | Seedling | 2021 | FNAMS – 49 | 6136518 | 59 | 5923 | 1 | | | | | | 1 | | PRJNA1041598 |
| 9021 | Sphingomonas | SAMN38288761 | Phaseolus vulgaris Vezer | Seedling | 2021 | FNAMS – 49 | 4314549 | 66 | 3872 | 0 | | | | | | | | PRJNA1041598 |
| 9023 | Microbacterium | SAMN38288762 | Phaseolus vulgaris Flavert | Seedling | 2021 | FNAMS – 49 | 3657683 | 68 | 3530 | 0 | | | | | | | | PRJNA1041598 |
| 9026 | Massilia | SAMN38288763 | Phaseolus vulgaris Linex | Seedling | 2021 | FNAMS – 49 | 5960194 | 65 | 5128 | 2 | | | 1 | | 1 | | | PRJNA1041598 |
| 9029 | Frigoribacterium | SAMN38288769 | Phaseolus vulgaris Flavert | Seedling | 2021 | FNAMS – 49 | 3208820 | 71 | 2943 | 0 | | | | | | | | PRJNA1041598 |
| 9030 | Frigoribacterium | SAMN38288764 | Phaseolus vulgaris Deezer | Seedling | 2021 | FNAMS – 49 | 3266158 | 71 | 3014 | 0 | | | | | | | | PRJNA1041598 |
| 9031 | Flavobacterium | SAMN38288765 | Phaseolus vulgaris Vanilla | Seedling | 2021 | FNAMS – 49 | 5602649 | 34 | 4794 | 1 | | | | | | | 1 | PRJNA1041598 |
| 9032 | Pedobacter | SAMN38288766 | Phaseolus vulgaris Vanilla | Seedling | 2021 | FNAMS – 49 | 5582692 | 38 | 4687 | 1 | | | | | | | 1 | PRJNA1041598 |
| 9033 | Rathayibacter festucae | SAMN38288767 | Phaseolus vulgaris Linex | Seedling | 2021 | FNAMS – 32 | 4341678 | 72 | 3923 | 0 | | | | | | | | PRJNA1041598 |
| 9034 | Microbacterium | SAMN38288768 | Phaseolus vulgaris Vanilla | Seed | 2021 | FNAMS – 49 | 3691531 | 70 | 3354 | 0 | | | | | | | | PRJNA1041598 |
| 13502 | Pseudomonas fluorescens subgroup | SAMN09044009 | Raphanus sativus Flamboyant5 | Seed | 2013 | FNAMS – 49 | 6201658 | 61 | 5582 | 1 | | | 1 | | | | | PRJNA454856 |
| 13503 | Stenotrophomonas rhizophila | SAMN09062466 | Raphanus sativus Flamboyant5 | Seed | 2013 | FNAMS – 49 | 4875264 | 67 | 4288 | 1 | | | | | 1 | | | PRJNA459523 |
| 13504 | Pseudomonas koreensis subgroup | SAMN09062476 | Raphanus sativus Flamboyant5 | Seed | 2013 | FNAMS – 49 | 6595509 | 59 | 5839 | 3 | 2 | | | 1 | | | | PRJNA459522 |
| 13505 | Pantoea agglomerans | SAMN09062489 | Raphanus sativus Flamboyant5 | Seed | 2014 | FNAMS – 49 | 4932464 | 55 | 4512 | 1 | | | 1 | | | | | PRJNA459521 |
| 13506 | Pseudomonas fluorescens subgroup | SAMN09062490 | Raphanus sativus Flamboyant5 | Seed | 2014 | FNAMS – 49 | 5914838 | 61 | 5354 | 2 | 1 | | 1 | | | | | PRJNA459520 |
| 13507 | Pseudomonas viridiflava | SAMN09062491 | Raphanus sativus Flamboyant5 | Seed | 2014 | FNAMS – 49 | 5925962 | 59 | 5232 | 2 | 1 | | 1 | | | | | PRJNA459519 |
| 13508 | Pseudomonas koreensis subgroup | SAMN09062492 | Raphanus sativus Flamboyant5 | Seed | 2014 | FNAMS – 49 | 6031678 | 60 | 5258 | 1 | 1 | | | | | | | PRJNA459518 |
| 13509 | Pseudomonas fluorescens subgroup | SAMN09062824 | Raphanus sativus Flamboyant5 | Seed | 2014 | FNAMS – 49 | 6834697 | 61 | 6091 | 2 | 1 | | 1 | | | | | PRJNA459517 |
| 13510 | Pseudomonas fluorescens subgroup | SAMN09062823 | Raphanus sativus Flamboyant5 | Seed | 2014 | FNAMS – 49 | 6347968 | 60 | 5627 | 2 | 1 | | 1 | | | | | PRJNA459516 |
| 13511 | Erw inia persicina | SAMN09063090 | Raphanus sativus Flamboyant5 | Seed | 2014 | FNAMS – 49 | 5203109 | 55 | 4848 | 2 | | 2 | | | | | | PRJNA459515 |
| 13512 | Paenibacillus | SAMN09063421 | Raphanus sativus Flamboyant5 | Seed | 2014 | FNAMS – 49 | 5251198 | 39 | 4622 | 0 | | | | | | | | PRJNA459514 |
| 13513 | Plantibacter | SAMN09063410 | Raphanus sativus Flamboyant5 | Seed | 2014 | FNAMS – 49 | 4231965 | 69 | 3901 | 0 | | | | | | | | PRJNA459513 |
| 13514 | Pseudomonas fluorescens subgroup | SAMN09063676 | Raphanus sativus Flamboyant5 | Seed | 2014 | FNAMS – 49 | 6238853 | 60 | 5614 | 2 | 1 | | 1 | | | | | PRJNA459512 |
| 13515 | Pseudomonas viridiflava | SAMN09064882 | Raphanus sativus Flamboyant5 | Seed | 2014 | FNAMS – 49 | 5869336 | 59 | 5208 | 2 | 1 | | 1 | | | | | PRJNA459504 |
| 13516 | Pantoea agglomerans | SAMN09064883 | Raphanus sativus Flamboyant5 | Seed | 2014 | FNAMS – 49 | 5093661 | 55 | 4698 | 2 | | 1 | 1 | | | | | PRJNA459503 |
| 13517 | Pseudomonas fluorescens subgroup | SAMN09064884 | Raphanus sativus Flamboyant5 | Seed | 2014 | FNAMS – 49 | 5954142 | 60 | 5358 | 2 | 1 | | 1 | | | | | PRJNA459502 |
| 13528 | Pseudomonas fluorescens subgroup | SAMN09064885 | Raphanus sativus Flamboyant5 | Seed | 2014 | FNAMS – 49 | 6043101 | 60 | 5311 | 3 | 1 | 1 | 1 | | | | | PRJNA459501 |
| 13529 | Stenotrophomonas rhizophila | SAMN09064919 | Raphanus sativus Flamboyant5 | Seed | 2014 | FNAMS – 49 | 4637306 | 67 | 3997 | 1 | | | | | 1 | | | PRJNA459500 |
| 13530 | Enterobacter cancerogenus | SAMN09064920 | Raphanus sativus Flamboyant5 | Seed | 2013 | FNAMS – 49 | 4901530 | 56 | 4559 | 3 | | 2 | 1 | | | | | PRJNA459499 |
| 13531 | Bacillus simplex | SAMN16237854 | Raphanus sativus Flamboyant5 | Seed | 2013 | FNAMS – 49 | 5965991 | 40 | 5792 | 0 | | | | | | | | PRJNA665156 |
| 13532 | Pantoea agglomerans | SAMN09064921 | Raphanus sativus Flamboyant5 | Seed | 2013 | FNAMS – 49 | 4934344 | 55 | 4597 | 2 | | 1 | 1 | | | | | PRJNA459498 |
| 13533 | Pseudomonas orientalis | SAMN16237857 | Raphanus sativus Flamboyant5 | Seed | 2013 | FNAMS – 49 | 6107913 | 60 | 5416 | 3 | 1 | 1 | 1 | | | | | PRJNA665156 |
| 13566 | Pantoea agglomerans | SAMN16237858 | Raphanus sativus Flamboyant5 | Seed | 2014 | FNAMS – 49 | 5123550 | 55 | 4807 | 2 | | 1 | 1 | | | | | PRJNA665156 |
| 13569 | Pantoea agglomerans | SAMN16237903 | Raphanus sativus Flamboyant5 | Seed | 2015 | NA | 4797171 | 55 | 4436 | 2 | | 1 | 1 | | | | | PRJNA665156 |
| 13570 | Plantibacter | SAMN16237824 | Raphanus sativus Flamboyant5 | Seed | 2013 | FNAMS – 49 | 4149054 | 69 | 3870 | 0 | | | | | | | | PRJNA665156 |
| 13571 | Pseudomonas syringae group | SAMN16237876 | Raphanus sativus Flamboyant5 | Seed | 2014 | FNAMS – 49 | 6026070 | 62 | 5339 | 2 | 1 | | 1 | | | | | PRJNA665156 |
| 13572 | Pseudomonas viridiflava | SAMN16237826 | Raphanus sativus Flamboyant5 | Seed | 2013 | FNAMS – 49 | 6020391 | 59 | 5349 | 3 | 1 | 1 | | | 1 | | | PRJNA665156 |
| 13573 | Pseudomonas fluorescens subgroup | SAMN16237813 | Raphanus sativus Flamboyant5 | Seed | 2013 | FNAMS – 49 | 6390377 | 60 | 5665 | 3 | 2 | | 1 | | | | | PRJNA665156 |
| 13574 | Pseudomonas syringae group | SAMN16237900 | Raphanus sativus Flamboyant5 | Seed | 2015 | NA | 6002142 | 62 | 5323 | | 1 | | 1 | | | | | PRJNA665156 |
| 13575 | Pseudomonas colepterorum | SAMN16237901 | Raphanus sativus Flamboyant5 | Seed | 2015 | NA | 4805492 | 62 | 4267 | 0 | | | | | | | | PRJNA665156 |
| 13578 | Pseudomonas viridiflava | SAMN16237030 | Phaseolus vulgaris Crosser (K79184) | Seed | NA | NA | 6147017 | 59 | 5501 | 3 | 2 | 1 | | | | | | PRJNA665156 |
| 13579 | Pantoea agglomerans | NA | Phaseolus vulgaris Linex (K79238) | Seed | NA | NA | 4832156 | 55 | 4462 | 2 | | 1 | 1 | | | | | #N/A |
| 13583 | Pantoea agglomerans | SAMN16237032 | Phaseolus vulgaris Crosser (K79184) | Seed | NA | NA | 4776328 | 55 | 4353 | 2 | | 1 | 1 | | | | | PRJNA665156 |
| 13593 | Pantoea agglomerans | SAMN16237033 | Phaseolus vulgaris Flavert (K79240) | Seed | NA | NA | 5130088 | 55 | 4736 | 3 | | 1 | 2 | | | | | PRJNA665156 |
| 13594 | Paenibacillus | SAMN16237029 | Phaseolus vulgaris Linex (K79238) | Seed | NA | NA | 7325832 | 46 | 6437 | 0 | | | | | | | | PRJNA665156 |
| 13595 | Bacillus megaterium | SAMN16237039 | Phaseolus vulgaris Flavert (K79240) | Seed | NA | NA | 5775775 | 38 | 5892 | 0 | | | | | | | | PRJNA665156 |
| 13596 | Bacillus megaterium | SAMN16236979 | Phaseolus vulgaris Rocdor (K79265) | Seed | NA | NA | 6069910 | 37 | 6201 | 0 | | | | | | | | PRJNA665156 |
| 13597 | Bacillus | SAMN16237031 | Phaseolus vulgaris Linex (K79238) | Seed | NA | NA | 5441368 | 41 | 5220 | 0 | | | | | | | | PRJNA665156 |
| 13599 | Pseudomonas colepterorum | SAMN16237062 | Phaseolus vulgaris Flavert | Seed | 2016 | FNAMS – 49 | 4843938 | 62 | 4277 | 0 | | | | | | | | PRJNA665156 |
| 13600 | Pantoea agglomerans | SAMN16236980 | Phaseolus vulgaris Rocdor (K79265) | Seed | NA | NA | 4885829 | 55 | 4470 | 2 | | 1 | 1 | | | | | PRJNA665156 |
| 13601 | Pseudomonas syringae group | SAMN16237043 | Phaseolus vulgaris Flavert | Flow er | 2016 | FNAMS – 49 | 6149110 | 62 | 5491 | 2 | 1 | | 1 | | | | | PRJNA665156 |
| 13602 | Pseudomonas putida group | SAMN16236977 | Phaseolus vulgaris Rocdor (K79265) | Seed | NA | NA | 4779684 | 62 | 4251 | 0 | | | | | | | | PRJNA665156 |
| 13603 | Sphingomonas | SAMN16237060 | Phaseolus vulgaris Flavert | Seed | 2016 | FNAMS – 49 | 4395607 | 65 | 3939 | 0 | | | | | | | | PRJNA665156 |
| 13604 | Pseudomonas fluorescens subgroup | SAMN16237065 | Phaseolus vulgaris Flavert | Seed | 2016 | FNAMS – 49 | 6242560 | 60 | 5614 | 2 | 1 | | 1 | | | | | PRJNA665156 |
| 13605 | Frigoribacterium | SAMN16237059 | Phaseolus vulgaris Flavert | Seed | 2016 | FNAMS – 49 | 3326223 | 71 | 3094 | 0 | | | | | | | | PRJNA665156 |
| 13611 | Pseudomonas fluorescens subgroup | SAMN16237055 | Phaseolus vulgaris Flavert | Pods | 2016 | FNAMS – 49 | 6046369 | 60 | 5448 | 3 | 1 | 1 | 1 | | | | | PRJNA665156 |
| 13615 | Pseudomonas colepterorum | SAMN16237070 | Phaseolus vulgaris Flavert | Seeds | 2016 | FNAMS – 49 | 4842123 | 62 | 4282 | 0 | | | | | | | | PRJNA665156 |
| 13616 | Pantoea agglomerans | SAMN16237057 | Phaseolus vulgaris Flavert | Seeds | 2016 | FNAMS – 49 | 4780294 | 55 | 4376 | 2 | | 1 | 1 | | | | | PRJNA665156 |
| 13617 | Microbacterium | SAMN16237041 | Phaseolus vulgaris Flavert | Flow er | 2016 | FNAMS – 49 | 3559858 | 71 | 3281 | 0 | | | | | | | | PRJNA665156 |
| 13644 | Rhizobium | SAMN16237137 | Brassica napus Aviso | Seed | 2017 | IGEPP | 5086082 | 57 | 4731 | 1 | | | | | | 1 | | PRJNA665156 |
| 13647 | Massilia | SAMN16237783 | Brassica napus Tenor | Seed | 2017 | IGEPP | 6073959 | 65 | 5126 | 2 | | | 1 | | 1 | | | PRJNA665156 |
| 13649 | Pseudomonas fluorescens subgroup | SAMN16237913 | Raphanus sativus Flamboyant5 | Flow er | 2016 | FNAMS – 49 | 6315997 | 60 | 5711 | 3 | 1 | 1 | 1 | | | | | PRJNA665156 |
| 13700 | Pseudomonas fluorescens subgroup | SAMN16237051 | Phaseolus vulgaris Flavert | Flow er | 2016 | FNAMS – 49 | 6020185 | 60 | 5337 | 3 | 1 | 1 | 1 | | | | | PRJNA665156 |
| 13701 | Pseudomonas orientalis | SAMN16237071 | Phaseolus vulgaris Flavert | Seeds | 2016 | FNAMS – 49 | 5766540 | 60 | 5067 | 2 | 1 | | 1 | | | | | PRJNA665156 |
| 13702 | Pseudomonas fluorescens subgroup | SAMN16237046 | Phaseolus vulgaris Flavert | Flow er | 2016 | FNAMS – 49 | 6019882 | 60 | 5300 | 3 | 1 | 1 | 1 | | | | | PRJNA665156 |
| 13703 | Pseudomonas orientalis | SAMN16237069 | Phaseolus vulgaris Flavert | Seed | 2016 | FNAMS – 49 | 5804470 | 60 | 5113 | 2 | 1 | | 1 | | | | | PRJNA665156 |
| 13706 | Sphingomonas | SAMN16237122 | Brassica napus Astrid | Seed | 2017 | IGEPP | 4753019 | 66 | 4265 | 0 | | | | | | | | PRJNA665156 |
| 13707 | Frigoribacterium | SAMN16237123 | Brassica napus Astrid | Seed | 2017 | IGEPP | 3554678 | 73 | 3269 | 0 | | | | | | | | PRJNA665156 |
| 13708 | Oxalobacteraceae | SAMN16237124 | Brassica napus Astrid | Seed | 2017 | IGEPP | 5310716 | 64 | 4577 | 1 | | | | | 1 | | | PRJNA665156 |
| 13709 | Pantoea agglomerans | SAMN16237131 | Brassica napus Astrid | Seed | 2017 | IGEPP | 5248093 | 55 | 4996 | 2 | | 1 | 1 | | | | | PRJNA665156 |
| 13710 | Pseudomonas putida group | SAMN16237132 | Brassica napus Astrid | Seed | 2017 | IGEPP | 4671763 | 62 | 4146 | 0 | | | | | | | | PRJNA665156 |
| 13711 | Pseudomonas | SAMN16237133 | Brassica napus Aviso | Seed | 2017 | IGEPP | 6192813 | 60 | 5442 | 3 | 1 | 1 | | | 1 | | | PRJNA665156 |
| 13712 | Frigoribacterium | SAMN16237134 | Brassica napus Aviso | Seed | 2017 | IGEPP | 3155852 | 71 | 2911 | 0 | | | | | | | | PRJNA665156 |
| 13713 | Pseudomonas fluorescens subgroup | SAMN16237135 | Brassica napus Aviso | Seed | 2017 | IGEPP | 6833365 | 60 | 6129 | 5 | 2 | | 3 | | | | | PRJNA665156 |
| 13714 | Sphingomonas | SAMN16237136 | Brassica napus Aviso | Seed | 2017 | IGEPP | 4419935 | 66 | 3950 | 0 | | | | | | | | PRJNA665156 |
| 13715 | Pseudomonas | SAMN16237138 | Brassica napus Aviso | Seed | 2017 | IGEPP | 6196569 | 60 | 5438 | 3 | 1 | 1 | | | 1 | | | PRJNA665156 |
| 13716 | Pseudomonas fluorescens subgroup | SAMN16237140 | Brassica napus Aviso | Seed | 2017 | IGEPP | 7815059 | 59 | 7179 | 5 | 2 | | 3 | | | | | PRJNA665156 |
| 13717 | Rhizobium | SAMN16237141 | Brassica napus Aviso | Seed | 2017 | IGEPP | 5158824 | 57 | 4777 | 1 | | | | | | 1 | | PRJNA665156 |
| 13718 | Stenotrophomonas | SAMN16237142 | Brassica napus Aviso | Seed | 2017 | IGEPP | 4202066 | 66 | 3684 | 0 | | | | | | | | PRJNA665156 |
| 13719 | Pseudomonas putida group | SAMN16237175 | Brassica napus Boston | Seed | 2017 | IGEPP | 5697150 | 61 | 5297 | 1 | 1 | | | | | | | PRJNA665156 |
| 13720 | Sphingomonas | SAMN16237345 | Brassica napus Boston | Seed | 2017 | IGEPP | 3692396 | 66 | 3327 | 0 | | | | | | | | PRJNA665156 |
| 13721 | Massilia | SAMN16237342 | Brassica napus Boston | Seed | 2017 | IGEPP | 6071750 | 65 | 5129 | 2 | | | 1 | | 1 | | | PRJNA665156 |
| 13722 | Pseudomonas fluorescens subgroup | SAMN16237190 | Brassica napus Boston | Seed | 2017 | IGEPP | 6208084 | 61 | 5653 | 2 | 1 | | 1 | | | | | PRJNA665156 |
| 13723 | Pseudomonas lutea group | SAMN16237352 | Brassica napus Boston | Seed | 2017 | IGEPP | 5760407 | 60 | 5089 | 3 | 1 | 1 | 1 | | | | | PRJNA665156 |
| 13724 | Stenotrophomonas | SAMN16237355 | Brassica napus Boston | Seed | 2017 | IGEPP | 3959299 | 67 | 3423 | 0 | | | | | | | | PRJNA665156 |
| 13725 | Stenotrophomonas | SAMN16237356 | Brassica napus Colvert | Seed | 2017 | IGEPP | 4180738 | 66 | 3656 | 0 | | | | | | | | PRJNA665156 |
| 13726 | Rhizobium | SAMN16237357 | Brassica napus Colvert | Seed | 2017 | IGEPP | 4933550 | 60 | 4657 | 0 | | | | | | | | PRJNA665156 |
| 13727 | Pseudomonas putida group | SAMN16237372 | Brassica napus Colvert | Seed | 2017 | IGEPP | 4608063 | 62 | 4090 | 0 | | | | | | | | PRJNA665156 |
| 13728 | Sphingomonas | SAMN16237374 | Brassica napus Colvert | Seed | 2017 | IGEPP | 4411320 | 65 | 4033 | 0 | | | | | | | | PRJNA665156 |
| 13729 | Frigoribacterium | SAMN16237389 | Brassica napus Colvert | Seed | 2017 | IGEPP | 3556539 | 72 | 3236 | 0 | | | | | | | | PRJNA665156 |
| 13730 | Oxalobacteraceae | SAMN16237464 | Brassica napus Colvert | Seed | 2017 | IGEPP | 5306600 | 64 | 4569 | 1 | | | | | 1 | | | PRJNA665156 |
| 13731 | Pantoea agglomerans | SAMN16237466 | Brassica napus Colvert | Seed | 2017 | IGEPP | 4984051 | 55 | 4623 | 2 | | 1 | 1 | | | | | PRJNA665156 |
| 13732 | Erw inia persicina | SAMN16237467 | Brassica napus Express | Seed | 2017 | IGEPP | 5183088 | 55 | 4835 | 3 | | 3 | | | | | | PRJNA665156 |
| 13733 | Sphingomonas | SAMN16237468 | Brassica napus Express | Seed | 2017 | IGEPP | 4645472 | 66 | 4117 | 0 | | | | | | | | PRJNA665156 |
| Xcc2 | Xanthomonas campestris pv. campestris | SAMN09068515 | Raphanus sativus Flamboyant5 | Seed | 2014 | FNAMS – 49 | 5161175 | 65 | 4358 | 0 | | | | | | | | PRJNA459480 |

### Table S2. Strains used to build the different SynComs in this study

| Strain | Taxonomy | Biosample | Host | Habitat | Freq (Simonin et al., 2022) | T6SS | i1 | i2 | i3 | i4b | i5 | SC1 | SC2 | SC3 | SC4 | SC5 | SCA | SCB | SC1_fitness (6h) | SC2_fitness (6h) | SC3_fitness (6h) | SC4_fitness (6h) | SC5_fitness (6h) | SCA_fitness (6h) | SCB_fitness (6h) | ratio CFU (6h) | Phenotype (6h) | ratio CFU (24h) | Phenotype (24h) |
|---|---|---|---|---|---|---|---|---|---|---|---|---|---|---|---|---|---|---|---|---|---|---|---|---|---|---|---|---|---|
| CFBP13507 | Pseudomonas viridiflava | SAMN09062491 | Raphanus sativus Flamboyant5 | Seed | 0,70 | 2 | 1 | - | 1 | - | - | 1 | - | - | - | - | - | - | 0,03 | - | - | - | - | - | - | - | - | - | - |
| CFBP13647 | Massilia | SAMN16237783 | B.napus Tenor | Seed | 0,01 | 2 | - | - | 1 | 1 | - | 1 | - | - | - | - | - | - | 0,08 | - | - | - | - | - | - | -15,24 | S | -21,96 | S |
| CFBP13574 | Pseudomonas syringae group | SAMN16237900 | Raphanus sativus Flamboyant5 | Seed | 0,41 | 2 | 1 | - | 1 | - | - | 1 | - | - | - | - | - | - | 0,09 | - | - | - | - | - | - | - | - | - | - |
| CFBP13517 | Pseudomonas fluorescens subgroup | SAMN09064884 | Raphanus sativus Flamboyant5 | Seed | 0,38 | 2 | 1 | - | 1 | - | - | 1 | - | - | - | - | - | - | 0,11 | - | - | - | - | - | - | - | - | - | - |
| CFBP13722 | Pseudomonas fluorescens subgroup | SAMN16237190 | B.napus Boston | Seed | 0,19 | 2 | 1 | - | 1 | - | - | 1 | - | - | - | - | - | - | 0,13 | - | - | - | - | - | - | - | - | - | - |
| CFBP8761 | Oxalobacteraceae | SAMN16237762 | B.napus Tenor | Seed | 0,00 | 1 | - | - | - | 1 | - | 1 | - | - | - | - | - | - | 0,20 | - | - | - | - | - | - | -9,22 | S | -27,20 | S |
| CFBP13509 | Pseudomonas fluorescens subgroup | SAMN09062824 | Raphanus sativus Flamboyant5 | Seed | 0,19 | 2 | 1 | - | 1 | - | - | 1 | - | - | - | - | - | - | 0,27 | - | - | - | - | - | - | - | - | - | - |
| CFBP13514 | Pseudomonas fluorescens subgroup | SAMN09063676 | Raphanus sativus Flamboyant5 | Seed | 0,13 | 2 | 1 | - | 1 | - | - | 1 | - | - | - | - | - | - | 0,29 | - | - | - | - | - | - | - | - | - | - |
| Xcc8004 | Xanthomonas campestris pv. campestri | SAMN37207934 | Brassica oleracea var. botrytis | Leaf | 0,27 | 0 | - | - | - | - | - | 1 | 1 | 1 | 1 | 1 | - | - | 0,63 | 0,45 | nd | nd | nd | - | - | -7,07 | S | -12,75 | S |
| CFBP8763 | Oxalobacteraceae | SAMN16237778 | B.napus Tenor | Seed | 0,06 | 2 | - | - | - | 2 | - | 1 | - | - | - | - | - | - | 1,20 | - | - | - | - | - | - | -1,13 | S | -5,81 | S |
| CFBP13503 | Stenotrophomonas rhizophila | SAMN09062466 | Raphanus sativus Flamboyant5 | Seed | 0,36 | 1 | - | - | - | 1 | - | 1 | 1 | 1 | 1 | 1 | 1 | 1 | 2,68 | 2,67 | 30,47 | 12,38 | 2,30 | 25,48 | 8,35 | - | | - | - |
| CFBP13709 | Pantoea agglomerans | SAMN16237131 | B.napus Astrid | Seed | 0,68 | 2 | - | 1 | 1 | - | - | - | - | - | 1 | - | 1 | - | - | - | - | 0,07 | - | 0,37 | - | -7,16 | S | -3,27 | S |
| CFBP13510 | Pseudomonas fluorescens subgroup | SAMN09062823 | Raphanus sativus Flamboyant5 | Seed | 0,04 | 2 | 1 | - | 1 | - | - | - | 1 | - | - | - | 1 | - | - | 0,11 | - | - | - | 0,51 | - | -5,45 | S | -6,83 | S |
| CFBP13725 | Stenotrophomonas | SAMN16237356 | B.napus Colvert | Seed | 0,21 | 0 | - | - | - | - | - | - | 1 | - | - | - | 1 | - | - | 0,12 | - | - | - | 0,56 | - | -5,58 | S | -22,71 | S |
| CFBP8803 | Erwinia persicina | SAMN16237911 | Raphanus sativus Flamboyant5 | Flower | 0,30 | 2 | - | 2 | - | - | - | - | - | 1 | - | - | 1 | - | - | - | 0,26 | - | - | 0,57 | - | -6,98 | S | -3,17 | S |
| CFBP13508 | Pseudomonas koreensis subgroup | SAMN09062492 | Raphanus sativus Flamboyant5 | Seed | 0,06 | 1 | 1 | - | - | - | - | - | - | 1 | - | - | - | - | - | - | 0,14 | - | - | - | - | - | - | - | - |
| CFBP13505 | Pantoea agglomerans | SAMN09062489 | Raphanus sativus Flamboyant5 | Seed | 0,88 | 1 | - | - | 1 | - | - | - | - | 1 | - | - | - | 1 | - | - | 0,71 | - | - | - | 0,35 | -5,41 | S | -1,00 | S |
| CFBP8758 | Pseudomonas putida group | SAMN16237690 | B.napus Major | Seed | 0,34 | 0 | - | - | - | - | - | - | - | - | - | 1 | - | - | - | - | - | - | 0,10 | - | - | - | - | - | - |
| CFBP13512 | Paenibacillus | SAMN09063421 | Raphanus sativus Flamboyant5 | Seed | 0,38 | 0 | - | - | - | - | - | - | - | - | - | 1 | - | - | - | - | - | - | 0,29 | - | - | -6,72 | S | -15,31 | S |
| CFBP8752 | Rhizobium | SAMN16237344 | B.napus Boston | Seed | 0,00 | 0 | - | - | - | - | - | - | - | - | - | 1 | - | - | - | - | - | - | 1,63 | - | - | -0,55 | R | -3,06 | S |
| CFBP13513 | Plantibacter | SAMN09063410 | Raphanus sativus Flamboyant5 | Seed | 0,35 | 0 | - | - | - | - | - | - | - | - | - | 1 | - | - | - | - | - | - | 4,38 | - | - | -2,57 | S | -4,26 | S |
| CFBP13533 | Pseudomonas orientalis | SAMN16237857 | Raphanus sativus Flamboyant5 | Seed | 0,09 | 3 | 1 | 1 | 1 | - | - | - | - | - | 1 | - | - | - | - | - | - | 0,03 | - | - | - | - | - | - | - |
| CFBP13711 | Pseudomonas | SAMN16237133 | B.napus Aviso | Seed | 0,01 | 3 | 1 | 1 | - | 1 | - | - | - | - | 1 | - | - | - | - | - | - | 0,03 | - | - | - | - | - | - | - |
| CFBP13569 | Pantoea agglomerans | SAMN16237903 | Raphanus sativus Flamboyant5 | Seed | 0,56 | 2 | - | 1 | 1 | - | - | - | - | - | 1 | - | - | - | - | - | - | 0,91 | - | - | - | - | - | - | - |
| CFBP13531 | Bacillus simplex | SAMN16237854 | Raphanus sativus Flamboyant5 | Seed | 0,00 | 0 | - | - | - | - | - | - | - | - | 1 | - | - | 1 | - | - | - | 11,01 | - | - | 3,02 | nd | nd | -1,58 | R |
| CFBP13571 | Pseudomonas syringae group | SAMN16237876 | Raphanus sativus Flamboyant5 | Seed | 0,24 | 2 | 1 | - | 1 | - | - | - | - | 1 | - | - | - | - | - | - | 0,20 | - | - | - | - | - | - | - | - |
| CFBP13530 | Enterobacter cancerogenus | SAMN09064920 | Raphanus sativus Flamboyant5 | Seed | 0,07 | 3 | - | 2 | 1 | - | - | - | - | 1 | - | - | - | - | - | - | 0,44 | - | - | - | - | - | - | - | - |
| CFBP13719 | Pseudomonas putida group | SAMN16237175 | B.napus Boston | Seed | 0,07 | 1 | 1 | - | - | - | - | - | - | 1 | - | - | - | - | - | - | 0,82 | - | - | - | - | - | - | - | - |
| CFBP8753 | Oxalobacteraceae | SAMN16237354 | B.napus Boston | Seed | 0,01 | 1 | - | - | - | 1 | - | - | - | 1 | - | - | - | 1 | - | - | 2,22 | - | - | - | 1,50 | -3,34 | S | -6,80 | S |
| CFBP13644 | Rhizobium | SAMN16237137 | B.napus Aviso | Seed | 0,31 | 1 | - | - | - | - | 1 | - | - | 1 | - | - | - | 1 | - | - | 4,13 | - | - | - | 1,70 | -0,19 | R | -0,55 | S |
| CFBP13718 | Stenotrophomonas | SAMN16237142 | B.napus Aviso | Seed | 0,02 | 0 | - | - | - | - | - | - | 1 | - | - | - | - | - | - | 0,16 | - | - | - | - | - | - | - | - | - |
| CFBP8771 | Pseudomonas putida group | SAMN16237697 | B.napus Major | Seed | 0,06 | 1 | 1 | - | - | - | - | - | 1 | - | - | - | - | - | - | 0,38 | - | - | - | - | - | - | - | - | - |
| CFBP13575 | Pseudomonas coleopterorum | SAMN16237901 | Raphanus sativus Flamboyant5 | Seed | 0,14 | 0 | - | - | - | - | - | - | 1 | - | - | - | - | - | - | 0,50 | - | - | - | - | - | - | - | - | - |
| CFBP8773 | Pseudomonas putida group | SAMN16237754 | B.napus Mohican | Seed | 0,11 | 0 | - | - | - | - | - | - | 1 | - | - | - | - | - | - | 0,53 | - | - | - | - | - | - | - | - | - |
| CFBP13724 | Stenotrophomonas | SAMN16237355 | B.napus Boston | Seed | 0,12 | 0 | - | - | - | - | - | - | 1 | - | - | - | - | - | - | 0,57 | - | - | - | - | - | -6,33 | S | -28,01 | S |
| CFBP13726 | Rhizobium | SAMN16237357 | B.napus Colvert | Seed | 0,10 | 0 | - | - | - | - | - | - | 1 | - | - | - | - | - | - | 1,96 | - | - | - | - | - | -1,69 | R | -9,92 | S |
| CFBP8762 | Rhizobium | SAMN16237769 | B.napus Tenor | Seed | nd | 0 | - | - | - | - | - | - | 1 | - | - | - | - | - | - | 2,06 | - | - | - | - | - | -0,79 | R | -4,16 | S |
| CFBP13714 | Sphingomonas | SAMN16237136 | B.napus Aviso | Seed | 0,08 | 0 | - | - | - | - | - | - | - | - | - | 1 | - | - | - | - | - | - | - | - | - | - | - | - | - |
| CFBP13729 | Frigoribacterium | SAMN16237389 | B.napus Colvert | Seed | 0,19 | 0 | - | - | - | - | - | - | - | - | 1 | - | - | - | - | - | - | - | - | - | - | - | - | - | - |
| CFBP8751 | Frigoribacterium | SAMN16237779 | B.napus Tenor | Seed | 0,02 | 0 | - | - | - | - | - | - | - | - | - | 1 | - | - | - | - | - | - | - | - | - | - | - | - | - |
| CFBP8757 | Aeromicrobium | SAMN16237654 | B.napus Major | Seed | 0,01 | 0 | - | - | - | - | - | - | - | - | - | 1 | - | - | - | - | - | - | - | - | - | - | - | - | - |
| CFBP8759 | Frigoribacterium | SAMN16237713 | B.napus Mohican | Seed | 0,06 | 0 | - | - | - | - | - | - | - | - | 1 | - | - | - | - | - | - | - | - | - | - | - | - | - | - |
| CFBP13733 | Sphingomonas | SAMN16237468 | B.napus Express | Seed | 0,10 | 0 | - | - | - | - | - | - | - | 1 | - | - | - | - | - | - | nd | - | - | - | - | - | - | - | - |
| CFBP13708 | Oxalobacteraceae | SAMN16237124 | B.napus Astrid | Seed | 0,02 | 1 | - | - | - | 1 | - | - | - | - | 1 | - | - | - | - | - | - | nd | - | - | - | -6,67 | S | -21,29 | S |
| CFBP13720 | Sphingomonas | SAMN16237345 | B.napus Boston | Seed | 0,01 | 0 | - | - | - | - | - | - | - | - | 1 | - | - | - | - | - | - | nd | - | - | - | - | - | - | - |
| CFBP8764 | Sphingomonas | SAMN16237800 | B.napus Tenor | Seed | 0,09 | 0 | - | - | - | - | - | - | - | - | - | 1 | - | - | - | - | - | - | nd | - | - | - | - | - | - |
| CFBP8790 | Microbacterium | SAMN16238272 | Raphanus sativus Flamboyant5 | Seed | 0,07 | 0 | - | - | - | - | - | - | - | - | - | 1 | - | - | - | - | - | - | nd | - | - | -0,20 | R | -0,19 | R |

### Table S3. Percentage of variance in bacterial phylogenetic composition of SynCom 1 to 5 (SC1-SC5) explained by initial stains (Strain) composition, time of confrontation (6h and 24h, Time) or Strain x Time interaction.

| SynCom | Strain | Time | Strain x Time |
|---|---|---|---|
| SC1 | 36.9 | 30.8 | ns |
| SC2 | 29.2 | 37.4 | 24.8 |
| SC3 | 26.2 | 64.3 | 5.2 |
| SC4 | 38.9 | 35.1 | 18.1 |
| SC5 | 75.1 | 6.2 | 13.5 |
